# Using high-throughput barcode sequencing to efficiently map connectomes

**DOI:** 10.1101/099093

**Authors:** Ian D Peikon, Justus M Kebschull, Vasily V Vagin, Diana I Ravens, Yu-Chi Sun, Eric Brouzes, Ivan R Corrêa, Dario Bressan, Anthony Zador

**Affiliations:** Watson School of Biological Sciences, Cold Spring Harbor Laboratory, Cold Spring Harbor, New York 11724, USA; Cold Spring Harbor Laboratory, Cold Spring Harbor, New York 11724, USA; Department of Biomedical Engineering, Stony Brook University, Stony Brook, New York 11794, USA; Laufer Center for Physical and Quantitative Biology, Stony Brook University, Stony Brook, New York 11794, USA; New England Biolabs, Inc., Ipswich, Massachausetts 01938, USA; Cancer Research UK Cambridge Institute, Li Ka Shing Centre, University of Cambridge, Cambridge CB2 0RE, United Kingdom

## Abstract

The function of a neural circuit is determined by the details of its synaptic connections. At present, the only available method for determining a neural wiring diagram with single synapse precision—a “connectome”—is based on imaging methods that are slow, labor-intensive and expensive. Here we present SYNseq, a method for converting the connectome into a form that can exploit the speed and low cost of modern high-throughput DNA sequencing. In SYNseq, each neuron is labeled with a unique random nucleotide sequence—an RNA “barcode”—which is targeted to the synapse using engineered proteins. Barcodes in pre- and postsynaptic neurons are then associated through protein-protein crosslinking across the synapse, extracted from the tissue, and then joined into a form suitable for sequencing. Although at present the inefficiency in our hands of barcode joining precludes the widespread application of this approach, we expect that with further development SYNseq will enable tracing of complex circuits at high speed and low cost.

## Introduction

The brain is extraordinarily complex, consisting of myriad neurons connected by even larger numbers of synapses. Disruption of these connections contributes to many neuropsychiatric disorders including autism, schizophrenia and depression. Understanding how the brain processes information and produces actions requires knowledge of both the structure of neural circuits, and of the patterns of neural activity. Sophisticated technology for recording ever-larger numbers of neurons is now widely available and is providing unprecedented insight into the physiological responses of brain circuits [1,2]. In contrast, circuit-mapping technologies with synaptic resolution remain very slow, expensive and labor intensive.

Mapping neural connectivity is traditionally viewed as a problem of microscopy. Electron microscopy (EM) allows direct imaging of synaptic contacts between neurons, so in principle circuit mapping with EM is trivial. In practice, however, it is complicated by a mismatch of scales. Imaging synapses requires nanometer resolution. In contrast, brain circuits span macroscopic distances, from millimeters in small organisms to tens of centimeters in humans. Circuit reconstruction using EM thus needs to bridge these scales, resulting in the requirement that thin axonal processes be traced across thousands of sections at an exceedingly low error rate. For example, for a 5 mm axon, and EM sections 50 nm thick, the required accuracy per single axon section would need, under simple assumptions, to exceed 99.999 % in order to achieve a 36% chance of assigning a correct connection. Several major efforts are underway to increase the throughput and autonomy of EM and have resulted in impressive improvements of speed and scale [3–10]. Unfortunately, most of these advancements require very expensive instruments, and the challenge of automatically tracing of axonal processes through EM stacks remains unsolved.

Electrophysiological approaches allow probing the connectivity of pairs or small groups of nearby neurons [11–13]. These efforts have uncovered elements of high-order structure within neural circuits, as well as spatially intertwined but non-interconnected networks [12,14]. However, such physiological methods are labor-intensive, and cannot readily be scaled for the analysis of larger neural circuits or a full nervous system [15].

We have been developing high-throughput sequencing as a fast and efficient alternative to microscopy or physiology for probing neuroanatomical connectivity [16-18]. To translate anatomical questions to a format amenable to sequencing, we label neurons uniquely with random nucleic acid sequences (“barcodes”). As a first proof of principle, we recently described MAPseq, a method for reading out long range projections with single neuron resolution [18]. In MAPseq, we infect neurons with a pool of barcoded virus particles and thus uniquely label every infected neuron with the barcode sequence carried by the viral particle that infected the neuron. The barcode is then expressed as an mRNA and is transported into axons, where we detect the barcode mRNA by sequencing as a proxy for the axonal projection of every labeled neuron. MAPseq allows the simultaneous tracing of thousands and potentially millions of single neuron projections – presenting a speedup of up to five orders of magnitude over traditional, microscopy-based methods. While MAPseq provides information about area-to-area connectivity at single neuron resolution, it does not provide single-neuron information about neuron-to-neuron connectivity.

Here, we introduce SYNseq, a method for converting synaptic connections into a form suitable for readout by high-throughput DNA sequencing. SYNseq consists of four steps: neuronal barcoding, trafficking of barcodes to the synapses via tight association with engineered synaptic proteins, joining of barcodes into a form suitable for sequencing, and reconstruction of the network connectivity (Fig. 1). Briefly, a pre-synaptic mRNA barcode is trafficked to the presynaptic terminal via association with an engineered version of the Neurexin1B (Nrx1B) protein. Likewise, the postsynaptic barcode is trafficked to the postsynaptic terminal via association with a modified Neuroligin1AB (Nlg1AB) protein. Across a synapse, the presynaptic SYNseq components are covalently linked to the postsynaptic SYNseq components and then immunoprecipitated for further biochemical manipulation to link the pre- and postsynaptic barcodes.

**Figure 1.**
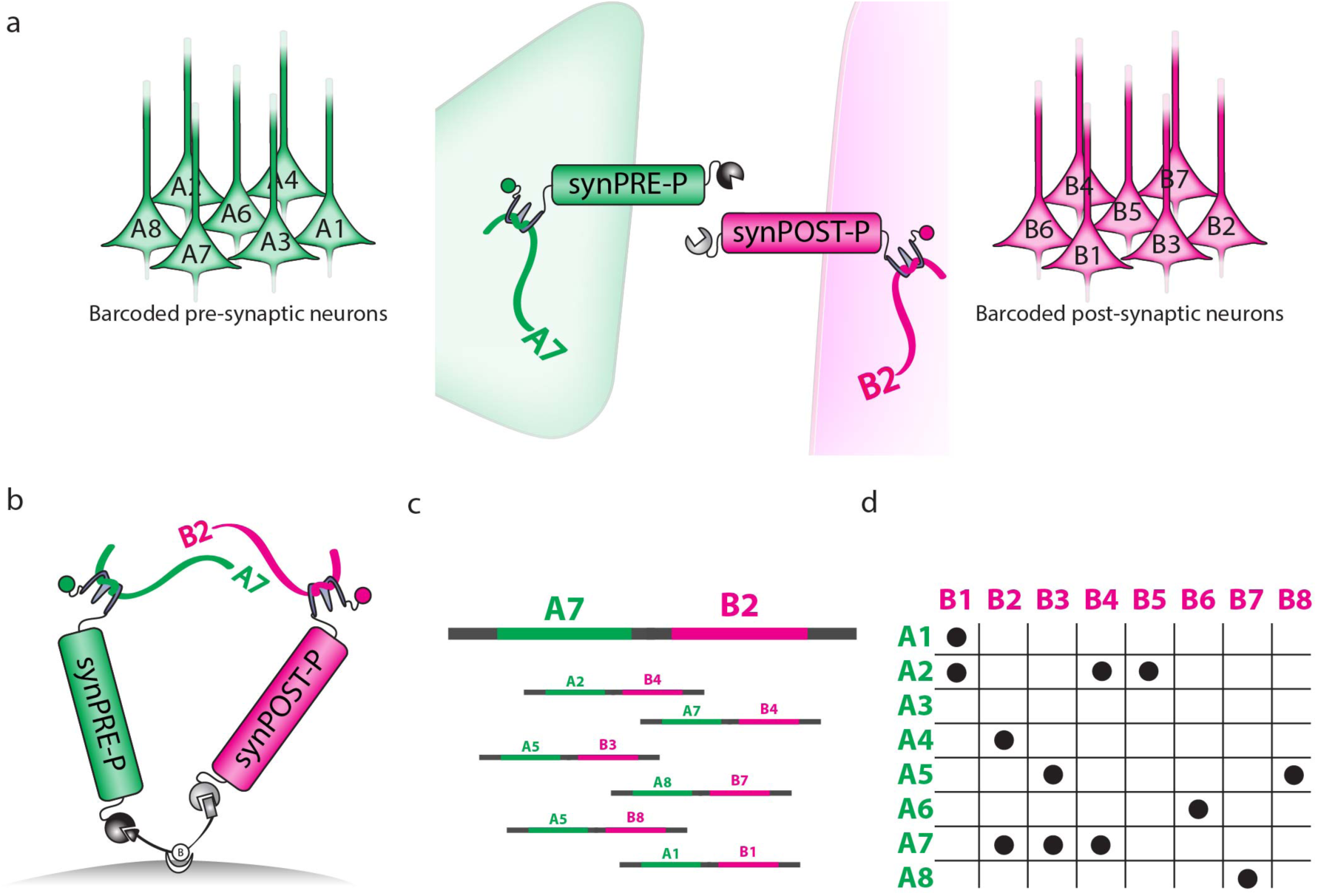
Overview of SYNseq. (a) First, we express random mRNA barcodes and modified synaptic proteins in a separate population of pre- and postsynaptic neurons, so that each neuron is uniquely labeled with a different barcode sequence (compare ref [18]). The modified synaptic proteins specifically bind the mRNA barcodes via an RNA binding domain, thereby trafficking barcodes to the pre- (*left*) or postsynaptic (*right*) compartments, respectively. The proteins meet at the synapse. (b) Next, the pre- and postsynaptic proteins are cross-linked *in situ* using a custom synthetic bivalent linker, which covalently joins the extracellular domains of the pre- and postsynaptic proteins. The resulting complexes, consisting of covalently bound pre- and postsynaptic protein pairs bound via the RNA binding domain to their respective barcodes, are then purified via immunoprecipitation (IP). (c) The associated barcode pairs, which represent pairs of connected neurons, are joined, amplified, and sequenced. (d) Sequencing data then allows the reconstruction of a connectivity matrix.

We develop a set of reagents that allow for synaptic trafficking of barcodes, specific crosslinking of carrier proteins across synapses, and recovery and joining of the barcode pairs that define a specific synapse. Unfortunately, however, the efficiency of the last step – barcode joining – is in our hands insufficient to provide for reliable synapse recovery while avoiding false positives. Despite this shortcoming, our work lays the foundation for using high-throughput DNA sequencing for circuit reconstruction. Future improvements upon SYNseq have the potential to enable high-throughput connectomics, opening up many new avenues of research in neuroscience.

## Results

In what follows, we first describe the design of the pre- and postsynaptic proteins and demonstrate their function in HEK cell culture. We then demonstrate the synaptic localization of the proteins in primary neuronal culture and describe the isolation of the transsynaptic complex formed by the proteins and their associated RNA barcodes. Finally we describe our efforts to join the pre- and postsynaptic barcodes of each complex by emulsion RT-PCR.

### Neuronal barcodes

We have previously developed a Sindbis-based system for over-expression of neuronal barcodes, which was used to map projections (MAPseq). This work revealed that unique barcodes can be efficiently targeted to individual neurons, and amplified through the Sindbis virus system. The core of this expression system that each virus particle carries a unique unique RNA barcode, which is extensively replicated after infection by the machinery of the Sindbis virus – and is flanked by sequences that allow for efficient recovery and sequencing. In this work, we sought to extend the MAPseq system by developing proteins that could specifically target these barcodes to the pre- and post-synaptic density, and enabling recovery of barcode pairs across individual synapses.

### Protein design

In order to enable recovery of synaptic connectivity from neuronal barcodes, we developed a pair of SYNseq carrier proteins that conform to three requirements. First, we required that they traffic to the pre- and postsynaptic terminals, respectively. Second, we required that the pre- and postsynaptic components be in a form that allowed for synapse-specific covalent cross-linking. Finally, we required that they bind the barcode mRNA tightly and specifically.

To design proteins that fulfill these requirements, we started with the presynaptic protein Nrx1B and the postsynaptic protein Nlg1AB. Both are relatively simple, single pass membrane proteins that traffic naturally to the appropriate side of a mammalian synapse.

To enable these proteins to carry barcode mRNAs, we inserted the RNA binding domain nλ into the cytoplasmic tail of the two proteins. The nλ domain is a 22 amino acid peptide (derived from the λ_N_ protein of λ bacteriophage) that specifically and strongly binds to a particular 15 nt RNA sequence, termed boxB [19]. We also added the corresponding boxB sequence to the barcode mRNA, causing the association of SYNseq proteins and barcode mRNAs.

To permit crosslinking of the proteins across the synapse, we fused the self-labeling proteins CLIP and SNAP to the extracellular domains of Nrx1B and Nlg1AB, respectively. CLIP and SNAP are engineered versions of a DNA repair enzyme that specifically react and form a covalent bond with benzylcytosine and benzylguanine [20,21], respectively. To crosslink the CLIP and SNAP domains, we synthesized a bi-functional small molecule that contains benzylcytosine on one end, and benzylguanine on the other, and is thus capable of forming a covalent bond between our pre- and postsynaptic proteins. In addition, this crosslinker contains a biotin functional group that allows efficient immunoprecipitation (IP). A second version of the crosslinker further allows for elution from beads after IP for biotin via a cleavable disulfide bridge (Supplemental Fig. 1). The two versions of the crosslinker (BG-PEG-Biotin-PEG-BC and BG-PEG-(S-S)-Biotin-PEG-BC) behave largely identically (data not shown) and we used them interchangeably in this study, unless elution was necessary. Finally, we fused a Myc epitope tag to Nrx1B, and an HA epitope tag to Nlg1AB, to aid in biochemistry and/or visualization.

The design of proteins that fulfilled all of the above requirements necessitated considerable troubleshooting. We tested various positions, RNA binding domains, and linker structures before finding a pair of proteins in which our modifications to Nrx1B and Nlg1AB did not disrupt the proteins’ endogenous trafficking pattern (Supplemental Fig 2,3,4,5; Supplemental Table 1). For ease of use and fast turnaround, we performed initial troubleshooting and benchmarking of different constructs by expression and co-expression in HEK cell culture. We then tested promising proteins for the more stringent criteria of synaptic localization and RNA binding in neuronal culture.

### Barcode mRNA design

The barcode mRNAs necessary for SYNseq underwent few changes from the initial design. Both pre- and postsynaptic barcode RNA carry a random 30 nt barcode sequence in their 5’ UTR, and four copies of the boxB sequence, which allows tight association of the barcode RNA to the SYNseq proteins, in the 3’ UTR. Finally, we introduced the coding sequences for GFP and mCherry into the presynaptic and postsynaptic barcode mRNAs, respectively.

### SYNseq protein traffic to the membrane, can be crosslinked and bind barcode mRNA in HEK cell culture

After troubleshooting (Supplemental Table 1), we focused on two proteins, Myc-CLIP-Nrx1B-1xnλ (synPRE-P) and HA-SNAP-Nlg1AB-1xnλ (synPOST-P), for further testing. Each contained a single repeat of nλ surrounded by long flexible linker sequences in their cytoplasmic tail, in a position that was previously reported not to disrupt the proteins’ endogenous trafficking [22–25]. The matching barcode mRNAs encode for GFP and mCherry, respectively, contain a random 30 nt barcode sequence in their 5’ UTR, and have four repeats of the boxB stem loop in their 3’UTR (Fig. 2A, B).

**Figure 2.**
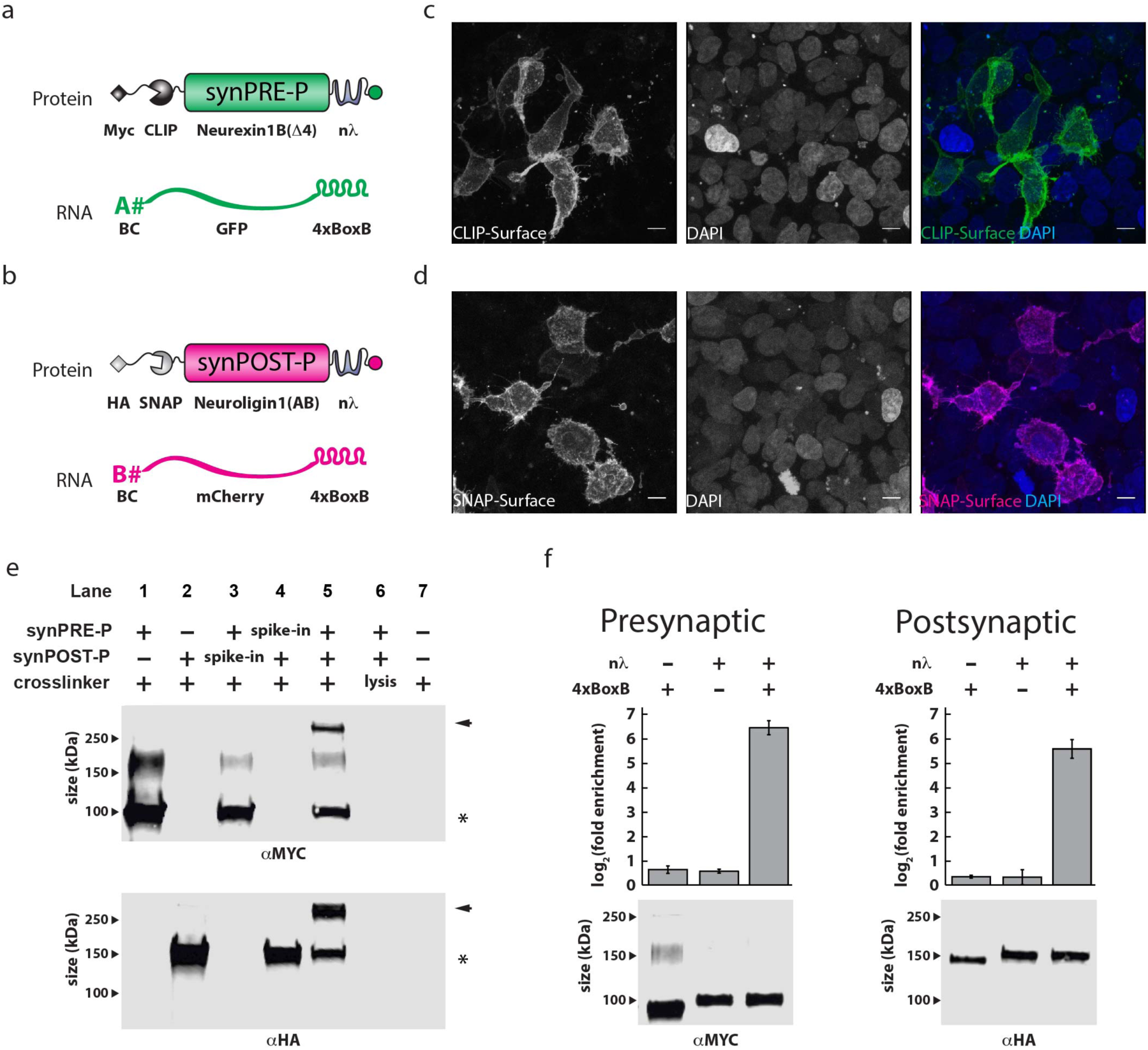
Optimization of SYNseq components in HEK cells. (a) The presynaptic components of the SYNseq system, consisting of CLIP-NRXN-1xnλ and a GFP encoding barcode RNA. (b) The postsynaptic components of SYNseq, consisting of SNAP-NLGN-1xnλ and a mCherry encoding barcode RNA. (c,d) A clear membrane staining of synPRE-P and synPOST-P can be observed after staining (c) synPRE-P expressing HEK cells with CLIP-Surface488 and (d) synPOST-P expressing HEK cells with SNAP-Surface488. Scale bar = 5 µm. (e) Western blot analysis shows that synPRE-P and synPOST-P can be specifically crosslinked by addition of a small molecule BG-PEG-Biotin-PEG-BC crosslinker. A crosslinked product is only produced when both synPRE-P and synPOST-P were expressed in HEK cells and the crosslinker was added before lysis (lane 5). Arrow = crosslinked band; star = uncrosslinked synPRE-P or POST. (f) synPRE-P and synPOST-P specifically and strongly bind to their respective barcode mRNAs as evident in RNA-IPs from transiently transfected HEK cells, after membrane tagging with BG-PEG-Biotin-PEG-BC. We show qRT-PCR analysis of 3 independent RNA-IP experiments and western blot analysis of a representative sample.

We first examined the localization of the two proteins. Both synPRE-P and synPOST-P traffic to the HEK cell membrane, as demonstrated by surface staining (Fig. 2C, D). This indicates that at least rudimentary trafficking of the proteins is uninterrupted by the addition of crosslinking and RNA binding domains.

We next tested whether we could specifically crosslink synPRE-P and synPOST-P in a simple scenario. We incubated HEK cells expressing synPRE-P and/or synPOST-P with the crosslinker, and performed an IP for the biotin on the crosslinker (Fig. 1B; Fig. 2E). We only detect a “crosslinked” synPRE-P—synPOST-P band (Fig. 2E arrow) in addition to the signals of the individual synPRE-P and synPOST-P proteins (Fig. 2E star), when both synPRE-P and synPOST-P proteins are expressed in the cells, and the crosslinker is added before cell lysis (Fig. 2E lane 5). Importantly, we do not detect this crosslinked band when only one of the two proteins is expressed (Fig. 2E lanes 1,2) or when the second protein was added post-lysis (Fig. 2E lanes 3,4). Moreover, the crosslinker is inactive in the lysis buffer (Fig. 2E lane 6) and does not result in nonspecific bands (Fig. 2E lane 7). These results indicate that we can form covalent synPRE-P—synPOST-P protein complexes, and that post-lysis association of synPRE-P and synPOST-P is low.

Finally, we confirmed that synPRE-P and synPOST-P specifically bind to barcode mRNA. After expression of either synPRE-P or synPOST-P in HEK cell culture and incubation with the biotinylated crosslinker, we find that barcode RNA is specifically enriched >32-fold by a pulldown for the biotinylated crosslinker (Fig. 2F). Importantly, the protein-RNA interaction depends on the presence of both the RNA binding domain nλ and its RNA recognition sequence boxB.

From these experiments, we confirm that synPRE-P and synPOST-P fulfill basic requirements for SYNseq proteins in HEK cell culture: membrane trafficking, efficient crosslinking and barcode mRNA binding. To further confirm these properties, we turned to more sensitive assays using neuronal cell culture.

### SYNseq proteins traffic to synapses in neuronal cell culture and can be crosslinked into transsynaptic complexes

Using neuronal cell culture, we next set out to characterize the behavior of synPRE-P and synPOST-P in neurons, including tests for synaptic localization and protein interaction using a series of increasingly stringent measures.

We first tested whether our protein modifications still permitted proper trafficking of each protein to the cell membrane in neurons. We used a double promoter Sindbis virus to express both the synPRE-P or synPOST-P protein and the pre- or postsynaptic barcode mRNA from a single virus (Fig. 3A, B), ensuring that SYNseq protein expression is always coupled with barcode mRNA expression. Sindbis virus is a positive strand RNA virus, characterized by a large (up to 6 kb [26]) payload as well as rapid and strong expression of transgenes. It therefore allows for rapid turnaround and iteration of constructs during troubleshooting. In addition, we have previously shown that Sindbis virus can be used to produce high diversity barcode libraries and can uniquely label neurons [18]. The Sindbis virus was thus well-suited to act as an expression vehicle for assessing SYNseq feasibility. Surface staining for Sindbis-expressed SYNseq proteins with CLIP-Surface488 (synPRE-P; Fig. 3C) and SNAP-Surface488 (synPOST-P; Fig. 3D) confirmed that our protein modifications did not interfere with proper trafficking of each protein to the cell membrane in neurons.

**Figure 3.**
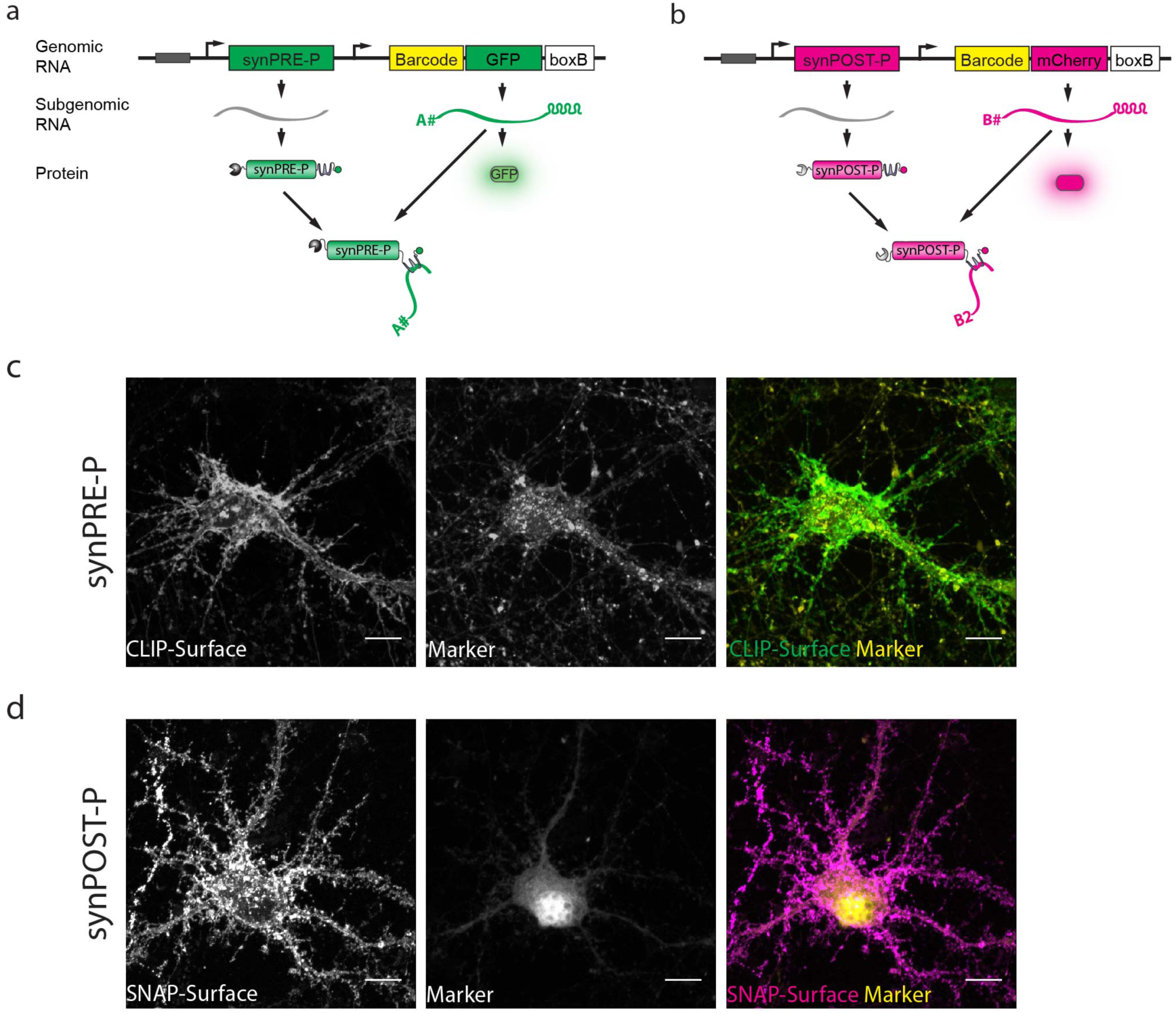
Viral expression and membrane trafficking of synPRE-P and synPOST-P in neurons. We use a double promoter Sindbis virus that expresses the (a) presynaptic or (b) postsynaptic components of SYNseq. (c,d) When expressed in neurons using Sindbis virus synPRE-P and synPOST-P show clear membrane trafficking as revealed by staining (c) synPRE-P with CLIP-Surface488 and (d) synPOST-P with SNAP-Surface488. Scale bar = 5 *µ*m.

We then used the proximity ligation assay (PLA) to screen for protein pairs that interact across synapses. PLA is a double antibody stain that results in a highly amplified signal only if the two probed epitopes are within 40 nm of each other (Fig. 4A, B, C, D) [27]. This interaction length is comparable to the dimensions of a synaptic cleft (20 nm [28]). The presence of a PLA signal between a pre- and a postsynaptic protein expressed in separate, but synaptically coupled populations of neurons thus provided a necessary but not sufficient criterion for establishing the synaptic localization of the two proteins (Fig. 4E, Supplemental Fig. 2,3,4,5, Supplemental Table 1). These experiments thus served as a screen for candidate protein pairs to be later tested in more stringent assays that restrict the interaction distance to < 5 nm.

**Figure 4.**
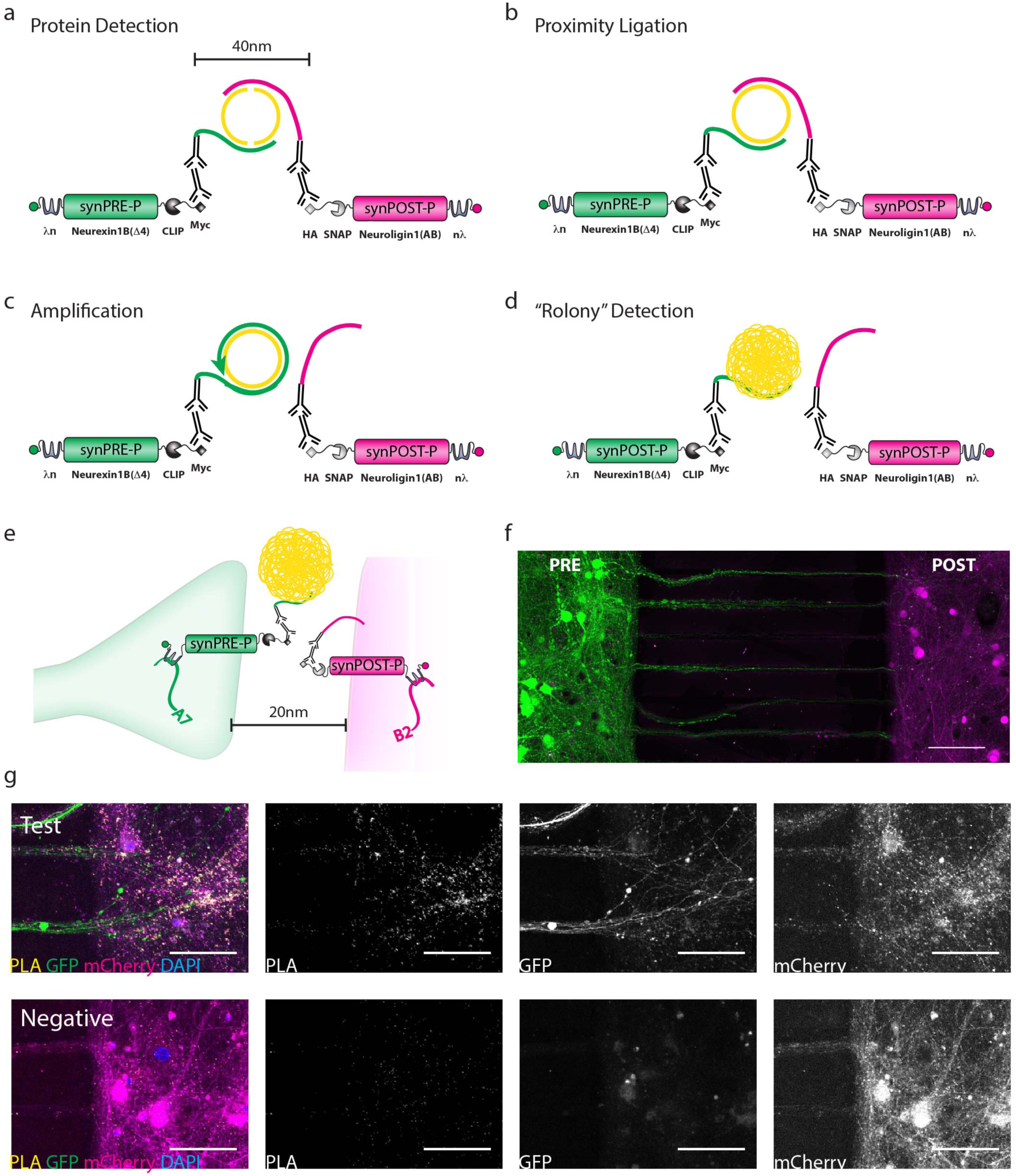
SynPRE-P and synPOST-P co-localize at synapses as measured by PLA. PLA is a multistep procedure involving (a) protein detection by two primary antibodies, followed by incubation with oligo-conjugated secondary antibodies. Two bridging oligos (yellow) are hybridized and will form a circle only if the two probed for epitopes are within 40 nm of each other. (b) The bridging oligos are ligated into a circle, which (c) is then amplified by rolling circle amplification and (d) detected by hybridizing fluorescently labeled oligos to the DNA ball. (e) If a labeled presynaptic cell forms a synapse with a labeled postsynaptic cell and synPRE-P and synPOST-P are co-localized at the synapse this should result in a PLA signal. (f) Two populations of neurons grown in a XONA microfluidic chamber system can be independently infected with Sindbis virus, as illustrated here by application of GFP-expressing virus on the left and mCherry expressing virus on the right. Scalebar = 100 *µ*m. (g) In this system PLA signals are clearly visible when synPRE-P positive (Gfp-labeled) axons come close to synPOST-P positive (mCherry-labeled) cells. The signal vastly exceeds the background staining observed in the absence of PRE. Scale bar = 50 *µ*m.

To achieve expression of synPRE-P and synPOST-P in separate, but synaptically coupled populations of neurons, we grew cultured primary hippocampal neurons in a XONA microfluidic device [29]. In a XONA, two chambers containing primary neurons are connected via long, thin microfluidic grooves. The setup ensures fluidic isolation of the two neural populations, allowing independent viral infection of the two populations, while permitting the growth of axons between chambers through the groves (Fig. 4F) [30].

We infected one population of cells with Sindbis virus expressing the pre-synaptic components and the other population of cells with barcoded Sindbis virus expressing the post-synaptic components. Using PLA we detected interaction between the synPRE-P and synPOST-P proteins preferentially where presynaptic (GFP positive) axons reach the postsynaptic chamber, suggesting that our proteins come within at least 40 nm of each other across separate but synaptically coupled cell populations (Fig. 4G).

PLA was a rapid and convenient screening tool to troubleshoot protein design (see Supplemental Fig. 2,3,4,5, Supplemental Table 1). However, its interaction distance of 40 nm was too large to confirm synaptic co-localization of synPRE-P and POST. In contrast, the bifunctional crosslinker designed to create the trans-synaptic synPRE-P—synPOST-P complex is only 4 nm long (Supplemental Fig. 1). Thus the creation of a synPRE-P—synPOST-P complex across synaptically coupled neuronal populations necessarily implies that the two proteins came within 4 nm of each other, satisfying even the strictest distance-based definitions of a synapse [4,31] (Fig. 5A). Again expressing synPRE-P and synPOST-P in synaptically coupled but separate neuronal populations using a XONA device, we found that we could specifically crosslink and immunoprecipitate synPRE-P and synPOST-P across the two populations of neurons, and by that logic, synapses (Fig. 5B). Note that, to simplify biochemistry from neuronal culture, we used a version of synPRE-P that contains a 3xFLAG epitope tag in addition to the myc epitope tag in this and all subsequent experiments.

**Figure 5.**
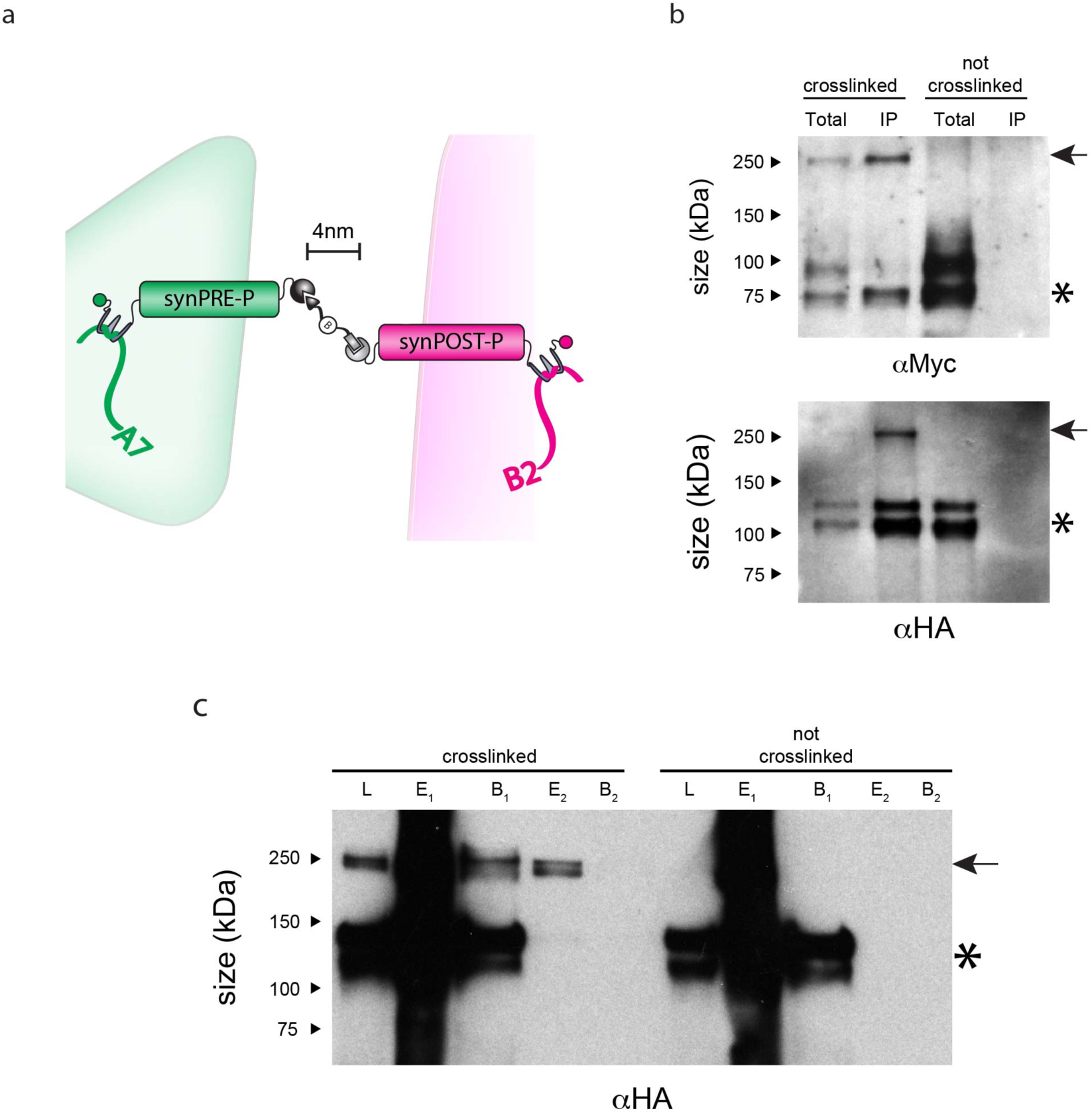
SynPRE-P and synPOST-P can be crosslinked into a transsynaptic complex and bind barcode mRNA. (a) If synPRE-P and synPOST-P interact at a synapse, we should be able to crosslink them with our 4 nm long crosslinker. (b) A synPRE-P—synPOST-P crosslinked band (arrow) can be observed from streptavidin IPs of separately infected XONA cultures only after addition of crosslinker. Un-crosslinked single proteins (star) can also be detected. (c) Performing two consecutive IPs, first for HA, then for 3xFLAG efficiently removes un-crosslinked proteins (star) and purifies synPRE-P—synPOST-P complexes (arrow). L = lysate; E1 = eluate off HA beads; B1 = remaining protein on HA beads; E2 = eluate off FLAG beads; B2 = remaining protein on HA beads.

### Enrichment for transsynaptic protein complexes in IP

When we immunoprecipitated crosslinked neuron samples from XONA cultures using the biotin tag on the crosslinker, we can detect at least three major products. First, we recover the desired synPRE-P—synPOST-P complex (Fig 5C, arrow) formed by interaction of the proteins across the synaptic cleft. However, we also observe signals arising from single synPRE-P and synPOST-P proteins that reacted with the one side of the crosslinker, but which have no partner on the other side of the crosslinker (Fig 5C, star). Such unpaired single proteins occur from at least three processes: from incomplete crosslinking across a synapse, from synapses at which only one side expresses a SYNseq protein, as well as from expression of either synPRE-P or synPOST-P in regions of the neuronal membrane that are not part of a synapse. Unfortunately, these unpaired proteins present a challenge to many barcode mRNA joining strategies. We therefore sought to remove them from the IP reaction by performing a double IP, selecting first for the postsynaptic HA tag, then for the presynaptic 3xFLAG tag, so that only proteins that contain both synPRE-P and synPOST-P remain after the second IP step. This strategy efficiently removed all unpaired synPRE-P or synPOST-P proteins from the IP and enriches for the transsynaptic protein complex (Fig. 5C).

From these results, we conclude that synPRE-P and synPOST-P conform to all three requirements that we had set out for SYNseq proteins: They specifically bind to barcode mRNA (Fig. 2F), traffic to synapses (Fig. 4, 5B) and can be specifically joined into a transsynaptic protein complex (Fig. 5B).

### Joining pre- and postsynaptic barcode mRNAs by emulsion RT-PCR

With the desired functional properties of synPRE-P and synPOST-P confirmed, we set out to develop a method for joining RNA barcodes specifically and efficiently. We tested a variety of methods (data not shown) before settling on barcode joining by emulsion reverse transcription-PCR (RT-PCR) followed by overlap PCR, which had previously proven successful for joining immunoglobulin mRNAs [32]. Briefly, we isolate synPRE-P—synPOST-P complexes by IP and emulsify the complexes using a microfluidic droplet system at a dilution such that each droplet contains no more than one complex. Inside each droplet, we then reverse transcribe the presynaptic and postsynaptic barcode mRNAs and subsequently join them into a single DNA strand by overlap PCR [32]. We then break the emulsion, isolate the barcode pairs and sequence them on an Illumina sequencing machine (Fig. 6A, Supplemental Fig. 6).

**Figure 6.**
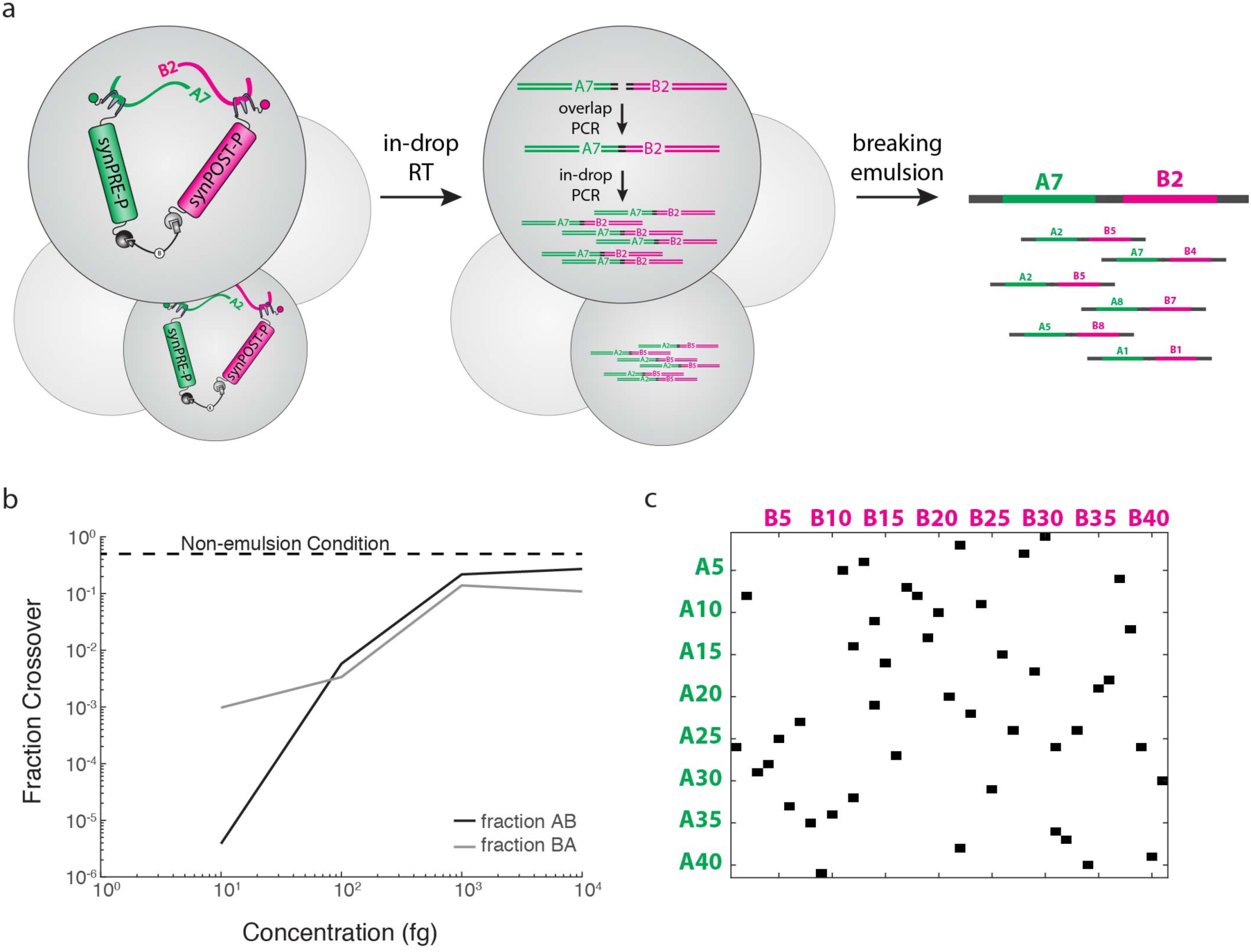
Emulsion RT-PCR can be used to join barcodes into barcode pairs. (a) Schematic of emulsion, in-drop RT and overlap PCR followed by breaking of the emulsion and purification of barcode pairs. (b) Dilution of input RNA decreases the false positive rate of emulsion PCR. Two separate barcode pairs (A-A and B-B) are subjected to uniform emulsions simultaneously to probe the concentrations at which overlap PCR becomes promiscuous. Fraction = 0.5 is equivalent to an aqueous (non-emulsified) PCR reaction. Here RNA molecules containing both a pre- and post- segment were employed. (c) “Connectivity matrix” obtained from emulsion RT-PCR followed by Illumina sequencing of synPRE-P—synPOST-P complexes from HEK cell culture, demonstrating that in principle SYNseq can be used to reconstruct synaptic circuits.

The success and failure of any method to join barcode mRNAs in SYNseq is measured not simply in the generation of barcode pairs, but critically in the ability of producing barcode pairs from mRNAs in the same complex, but not from mRNAs in different complexes or even unbound mRNAs. When using emulsion RT-PCR to join barcode mRNAs, such nonspecific joining occurs when more than one synPRE-P—synPOST-P complex, and thus more than one presynaptic or postsynaptic barcode mRNA are loaded into the same droplet. The fraction of unspecific joining events therefore ultimately depends on the concentration of complexes that are loaded into droplets and decreases with input concentration, as demonstrated by loading droplets with different concentrations of a test RNA that contains the sequences of both pre- and postsynaptic barcodes, but in inverted orientations relative to the final sequencing amplicon (Fig. 6B, Supplemental Fig. 7A).

To validate the emulsion RT-PCR approach, we first tested joining using protein-RNA complexes derived from HEK cell culture. To measure the rate of nonspecific joining of barcode mRNAs, we set up two experiments in parallel. Each experiment consists of one population of HEK cells that co-express synPRE-P, synPOST-P, and barcode mRNAs. In these experiments we used an earlier version of the SYNseq proteins. The barcode mRNAs in each experiment are tagged with an additional, experiment-specific sequence, a zip code (denoted here as A or B). We crosslinked and lysed each set of cells independently, and pooled the lysates. We then immunoprecipitated synPRE-P—synPOST-P complexes from the mixed lysate by streptavidin IP, released the complexes from the beads by cleaving the disulfide bridge on the crosslinker under mild reducing conditions (Supplemental Fig. 7B) and finally joined barcode mRNAs using emulsion RT-PCR. Sequencing the resulting barcode pairs revealed 42 presynaptic and 42 postsynaptic barcodes, which were joined into 45 unique barcode pairs (Fig. 6C).

Based on the experimental setup, barcode pairs should show matching zip codes (i.e. A-A pairs or B-B pairs), as those are the only biologically possible combinations. Any mixed barcode pairs (A-B or B-A) must arise from “crossover” events and therefore represent the rate of false-positives in recovery of the correct set of synaptic connections by SYNseq. In our HEK cell experiment we find a significant crossover rate of 13.3 % (barcodes from one experiment joined to barcodes of another experiment). Extrapolated to include undetectable A-A and B-B crossover events, this indicates an overall false-positive rate of 26.6 %. This false-positive rate, importantly, only measures false-positives arising from biochemical procedures post-lysis (i.e. protein-RNA dissociation, insufficient dilution for emulsions, etc.). It cannot quantify the false-positive barcode pairs that result from aberrant localization of the proteins or any other *in vivo* effects. The false-positive rate of this experiment appears to be dominated by the emulsion input concentration (Supplemental Fig. 7C), suggesting that we could reduce the false positive rate by further dilution of the complexes before emulsion.

We next tested emulsion RT-PCR for barcode joining in samples from neuronal cultures. We infected XONA devices with barcoded synPRE-P and synPOST-P Sindbis in the left and right chamber, respectively. Again, we set up two replicate experiments in parallel using experiment specific zip codes to differentiate the barcode mRNAs. We crosslinked, lysed and mixed the lysates from the two sets of XONA devices and performed a double immunoprecipitation to isolate synaptic synPRE-P—synPOST-P complexes. We then performed emulsion RT-PCR on a very dilute sample of the IP products to minimize the rate of crossover events.

Sequencing the resulting barcode pairs, we identified only 2 unique combinations of pre- and postsynaptic barcodes, albeit no false positive cross-experiment barcode pairs. While it is likely possible to increase this number of joined barcode pairs by increasing the number of droplets made, or by slightly increasing the concentration of emulsion input, we found ourselves in a regime far from the optimal necessary to achieve high throughput connectivity mapping by SYNseq. Taken together, our emulsion results suggest, that while SYNseq can be used to read out synaptic connectivity, barcode joining is currently a very low efficiency process, which will need to be optimized further before SYNseq can be used for circuit mapping.

## Discussion

We here present SYNseq, a scheme for converting synaptic connectivity into a form that can be deciphered using modern sequencing technology. SYNseq represents a significant step in the development of a method to map neuronal circuits with high throughput. We established each of the required biochemical components for SYNseq, namely barcoding [18], protein-joining, and RNA barcode joining (Fig. 1), but further work – in particular on barcode joining – will be needed to enable the robust tracing of neuronal circuits. We expect that an efficient and high-throughput method for determining both the local and long-range synaptic connectivity of neuronal populations would represent a transformative technology for research on neural circuits.

### DNA sequencing for neuroanatomy

Historically, the method of choice for inferring neuroanatomical connectivity has been microscopy. The specific form of microscopy used is determined by the resolution required to resolve the smallest important feature of the circuit in question. This implies that the smaller the structure of interest, the greater the resources and effort needed to perform the requisite imaging. Imaging methods often generate massive amounts of data, far beyond what is needed to answer simple questions about connectivity. These massive data sets can be challenging to store, and complicate analysis. Imaging therefore renders high-resolution neuroanatomic studies low throughput and limits the ability to scale to larger circuits. Because of these technical limitations, we currently have detailed circuit information about only a very limited number of mammalian circuits [6,33–35].

By applying modern high-throughput sequencing technology to neuroanatomical problems, we are attempting to overcome the imaging and tracing bottlenecks in high-resolution circuit analysis. DNA sequencing has undergone tremendous improvements in speed, throughput and cost in the last decade: While sequencing a human genome cost 3 billion dollars in 2000, it now costs less than 1000 dollars, and prices continue to fall at a rate exceeding Moore’s law [36,37]. The exceptional speed, parallelization and flexibility of modern sequencing technology thus motivate our attempts to translate the circuit features of interest into a format that is accessible to sequencing.

We recently presented MAPseq, the first application of DNA sequencing to neuroanatomy. MAPseq permits the simultaneous read out of thousands and potentially millions of single neuron long-range projections in a single mouse brain [18]. With SYNseq, we now provide a foundation for the considerably more challenging problem of determining synaptic connectivity. SYNseq takes inspiration from GRASP/mGRASP [38-40] and chromosome confirmation capture [41,42]. With further development, SYNseq may allow the routine mapping of complete circuits or even whole brains, on a time scale of weeks and at a cost measured in thousands rather than millions of dollars. The equipment needed for SYNseq is routinely available in many laboratories, and does not require major capital expenditures (with the ability to leverage existing core facilities and/or CROs for sequencing expertise). The high-throughput of SYNseq may make it possible to compare circuits across areas, species and treatments, and may lead to high-throughput screens for drugs or targets relevant to neurological disorders.

### Challenges ahead

Currently, the inefficiency of barcoding joining after pulldown limits the practical utility of SYNseq. In the present study we attempted to use emulsion overlap RT-PCR to join the pre- and postsynaptic RNAs held in one complex. Although we were able to show that this is technically possible (Fig. 6), we were unable to recover many barcode pairs after joining, rendering SYNseq in its current form unusable for answering biological questions. The most probable reason for our failure to join many barcodes is that single-molecule reverse transcription efficiency is low. Because reverse transcription must succeed for both the pre- and postsynaptic barcode RNA, each of which is only present in a single copy per loaded droplet, low RT efficiencies will have a devastating effect on the overall efficiency of barcode joining. Although we were unable to achieve efficient and specific joining of barcodes in droplets, it is possible that with further optimization a droplet-based strategy might succeed. Alternatively, joining strategies based on RNA ligation, gap filling or immobilization of complexes on beads, chips or in gels might emerge as more efficient strategies. Thus, although our present inability to achieve efficient and specific barcode joining limits the practical application of this approach, we expect the approach we have presented will form that foundation for an efficient method for deducing synaptic connections.

Another concern is that even brief (< 48 hour) expression of the synPRE-P and synPOST-P proteins, which are derived from synaptic proteins involved in synapses stabilization [43], could lead to formation of spurious synaptic connections. To circumvent this, the Nrx1B-Nlg1AB interaction domains could be eliminated [39], or similar approaches could be applied to other synaptic proteins that do not interact. Finally, all data presented here are based on using Sindbis virus to deliver barcodes and carrier proteins to neurons. Sindbis was a convenient choice because of its large (> 6 kb) payload, rapid (< 48 hr) expression and the ease of engineering. However, for some applications it may be advantageous to use a less toxic DNA virus amenable to cre-dependent control, such as adeno-assoicated virus (AAV) or herpes simplex virus (HSV) amplicons [44].

### Space and neuroanatomy

In its simplest incarnation, SYNseq fails to preserve the spatial relationships among neurons and their connections. Spatial resolution can, however, be recovered in a variety of ways, including dissection [16,18]. Most promisingly, emerging *in situ* sequencing methods developed by our lab and others (Chen *et al. in preparation*) [17,45] allow us to combine the advantages of high throughput sequencing with the spatial resolution required to ask many biological questions.

### Future directions

We envision that SYNseq will serve as a platform for future development towards encoding neural connectivity, and/or other biological variables – including cellular activity and/or identity – into a form that can be read by high-throughput sequencing, a technology which already operates at the scale of the complexity of neural circuits. Improvements to this method that allow for the co-expression of all of the components within the same cells – for example, by employing two separate RNA binding domains (Supplemental Fig. 5) – will allow for bi-directional and local-circuit mapping. Transgenic techniques, combined with an *in vivo* barcoding scheme based on recombinases [46–48] will allow for scaling this technology to mapping full brains.

High-throughput methods for mapping neuronal circuits will have a profound impact on neuroscience research. Circuits can be mapped at different developmental time-points, across animals with different genetic abnormalities, after discrete behavioral experiences, and/or after exposure to various pharmacological compounds. By employing DNA sequencing, the field of connectomics may enter an era of “big data” currently being experienced by other fields of biology, which were the first to harness DNA sequencing for deciphering genomes and transcriptomes.

## Experimental procedures

### Cell culture

We grew HEK293 cells under standard conditions at 37^o^C with 5 % CO_2_ in DMEM (Thermo Fisher) supplemented with 10 % heat inactivated FBS (Thermo Fisher) and 5 % Pen-Strep (Thermo Fisher). We cultured the cells in tissue culture plates coated with poly-L-lysine (Sigma-Aldrich) to aid adherence to the plate.

We prepared primary cultures of dissociated mouse hippocampal neurons from E18 mice as previously described [49]. Briefly, we isolated the brains and placed them into cold HBSS. We dissected hippocampi and incubated them in HBSS + trypsin + DNAse for 15 min at 37^o^C with gentle periodic agitation. We dissociated the cells and plated the cells at a density of 200,000 cells per well of a 12-well plate or 100,000 cells per side of a Xona Microfluidic Chamber (SND450, Xona Microfluidics). We cultured the neurons in Neurobasal (Thermo Fisher) + 2 % B27 (Thermo Fisher) + 1 % Pen-Strep (Thermo Fisher) + 0.5 mM L-glutamine (Thermo Fisher). For the first 4 days after dissociation, we added glutamate to the media at a final concentration of 125 μM. We incubated all neuronal cultures at 37^o^C with 5 % CO_2_ and conducted experiments after 14 days *in vitro*.

### Plasmid Construction

We constructed all plasmids used in this study by Gibson assembly [50] and standard cloning procedures and confirmed successful assembly by Sanger sequencing. All plasmids and annotated plasmid maps used in this study are available from Addgene. For accession numbers consult Supplemental Table 2.

We produced barcoded plasmid diversity libraries for both HEK cell transfection and Sindbis virus production as previously described [18] aiming for a minimum diversity of 10^6^ different plasmid sequences as estimated from colony counts.

### RNA Design

The presynaptic and postsynaptic RNA are based on GFP and mCherry coding sequences, respectively. In the 5’UTR of the preRNA we placed an anchor sequence (preHandle), a 100 bp spacer sequence, a NotI restriction site (for barcode cloning), the reverse complement of a qPCR tag denoting the library batch (A or B), a 30 nt barcode, the reverse complement sequence of the Illumina sequence P5-SBS3T, and an MluI restriction site for barcode cloning. In the 5’UTR of the postRNA we placed an anchor sequence (postHandle), a spacer sequence, an MluI restriction site (for BC cloning), the reverse complement of a qPCR tag denoting the library batch (A or B), a 30 nt barcode, the reverse complement sequence of the Illumina sequence P7-SBS8, and a NotI restriction site for barcode cloning. In the 3’UTR of both sequences, we placed four repeats of the boxB hairpin motif, 4xBoxB [19].

### Protein Design

We started with the interacting synaptic proteins – the presynaptic protein Neurexin1B (Uniprot Accesion Number: P0DI97, isoform 1b) and the post-synaptic protein Neuroligin1AB (Uniprot Accesion Number: Q99K10, isoform 1). In the final synPRE-P and synPOST-P proteins we fused MYC-CLIP [20] and HA-SNAP [21] to Nrx1B and Nlg1AB, respectively. In both cases, fusion was directly after the signal peptide sequence (after amino acid G46 in Nrx1B and K47 in Nlg1AB). We further fused a single copy of the nλ domain [19] flanked by flexible linkers after amino acid S423 in Nrx1B and after T776 in Nlg1AB.

### Proximity Ligation Assay

We fixed neurons grown in Xona microfluidic chambers in 4 % PFA in 0.1 M PBS for 15 min at room temperature, washed 2X in PBS, and quenched excess PFA by incubating 4 mM glycine in PBS for 5 min.

We then performed PLA according to the manufacturer’s instructions (Duolink PLA kit; Sigma-Aldrich). For the detection of the pre-synaptic proteins, we used a goat anti-MYC antibody ab9132 (Abcam), which recognizes the MYC tag on the extracellular domain of the presynaptic proteins. For detection of the post-synaptic proteins, we used a rabbit anti-HA antibody ab9110 (Abcam), which recognizes the HA tag on the extracellular domain of the postsynaptic proteins. We used Duolink secondary antibodies MINUS Probe Donkey anti-Rabbit (Sigma DUO92005) and PLUS Probe Donkey anti-Goat (Sigma DUO92003) and used the Duolink *in situ* far red detection reagents (Sigma DUO92013) for detection. All imaging was done using 20x objective on an LSM710 confocal microscope (Zeiss). One representative z-section is shown.

### Sindbis virus production and titering

We produced and titered all Sindbis viruses used in this study as previously described [49].

Briefly, we linearized the genomic plasmid and the DH(26S)5’SIN or DH-BB(5’SIN; TE12ORF) helper plasmid [49] by digestion with PacI or XhoI (New England Biolabs), respectively. We *in vitro* transcribed the linearized digestion products using the SP6 mMessage mMachine *in vitro* transcription system (Thermo Fisher) and transfected genomic and helper RNA into BHK cells using Lipofectamine 2000 (Thermo Fisher). We harvested virus 40 hrs after transfection and concentrated it by ultracentrifugation. We determined the titer of all viruses by qRT-PCR against a plasmid standard.

### Chemical syntheses of bifunctional crosslinkers

We prepared BG-PEG-Biotin-PEG-BC and BG-PEG-(S-S)-Biotin-PEG-BC by reacting BG-PEG-BC [51] with commercially available Biotin amidohexanoic acid NHS (Sigma-Aldrich) and NHS-S-S-dPEG^®^₄-biotin (Quanta Bio design), respectively. Briefly, we dissolved BG-PEG-BC (5.0 mg, 4.8 µmol) in anhydrous DMF (1.0 ml). We added biotin amidohexanoic acid NHS (2.3 mg, 5.0 µmol) or NHS-S-S-dPEG^®^₄-biotin (3.8 mg, 5.0 µmol), followed by triethylamine (1.0 µL, 7.2 µmol). We then stirred the reaction mixture overnight at room temperature. We removed the solvent under vacuum and purified the products by reversed-phase HPLC on a VYDAC 218TP series C18 column (22 x 250 mm, 10 µm particle size) at a flow rate of 20 ml/min using a water/acetonitrile gradient (0 to 95% acetonitrile over 45 min). Yields: BG-PEG-Biotin-PEG-BC (44%), ESI-MS *m/z* 1387.6916 (calc. for C_64_H_94_N_18_O_15_S^+^, *m/z* 1387.6940); BG-PEG-(S-S)-Biotin-PEG-BC (21%), ESI-MS *m/z* 1684.7621 (calc. for C_74_H_114_N_19_O_20_S_3_
^+^, *m/z* 1684.7644). We recorded high resolution mass spectra by electrospray ionization (ESI) on an Agilent 6210 Time-of-Flight (TOF) instrument.

### Tagging with BG/BC derivatives

We obtained all BG- and BC- functionalized derivatives, including the bifunctional cross-linkers (synthesis see above), CLIP-Surface488 and SNAP-Surface488 from New England Biolabs (NEB) and used them according to the manufacturer’s instructions. Briefly, we prepared a stock of 1 mM tag in DMSO and labeled cells expressing cell-surface CLIP-or SNAP-fusion proteins by incubating them in 5 μM tag in full media for 30 min, followed by three washes. We performed all imaging of surface labeled cells using a 63x objective on an LSM710 confocal microscope (Zeiss). 3D reconstructions of the images are shown to illustrate surface staining.

We slightly altered the above protocol for crosslinking cultured hippocampal neurons. Briefly, we incubated the cells with 2.5 μM bifunctional crosslinker in complete neuron media for 30 min. We then blocked all unreacted SNAP and CLIP epitopes by incubating the cells in 10 μM each of SNAP-cell block and CLIP-cell block (NEB S9106S and S9220S) in full neuron media for 30 min. We then washed the cells three times with full neuron media and finally in PBS.

### Protein and RNA IPs from HEK cells

After tagging with biotinylated linkers, we lysed cells in 1 ml of PLB3 lysis buffer (20 mM TrisHCl, 20 mM NaHCO_3_, 150 mM NaCl, 4 mM KCl, 2.5 mM MgCl_2_, 2.5 mM CaCl_2_, 10% glycerol, 1% CHAPS, 1% TritonX-100, 1x Complete EDTA-free protease inhibitor (Roche) [52]), per 12-well well of HEK cells or Xona microfluidic chamber. For RNA IPs we added RNase inhibitors RNAsin (Promega N2515) the PLB3 at 120 U/ml of lysis buffer.

We incubated the crude cell lysates on ice for 30 min, pelleted cell debris by centrifugation at 21,000 xg for 30 min at 4^o^C and saved a fraction of the resulting lysate for total protein analysis via western blot or other means. We loaded the remainder of the lysate onto 100 μl of pre-washed Dynabeads M280 Streptavidin magnetic beads (Thermo Fisher) and incubated with gentle rotation overnight at 4^o^C. We then washed the beads with PLB3 with ascending salt concentrations (2x with 150 mM NaCl, 2x with 300 mM NaCl, and 2x with 500 mM NaCl) for 10 min each.

We finally eluted the RNA or protein samples depending on the downstream application.

For emulsion RT-PCR we eluted protein samples from Dynabeads via addition of 15 mM DTT for 1 hr at 37^o^C.

For western blotting, we boiled the dynabeads in SDS containing loading buffer (New England Biolabs, B7709S) for 10 min.

For qPCR, we resuspended the beads in 400 μl Proteinase K buffer (100 mM Tris-HCL pH 7.5, 12.5 mM EDTA, 150 mM NaCl, 1 % SDS) with 2 μl of GlycoBlue (Thermo Fisher) and 20 μl of Proteinase K (20 μg/μl; Roche 03115879001). We incubated the samples at 65^o^C for 1 hr with gentle shaking and extracted the RNA using acid phenol: chloroform (Thermo Fisher) according to the manufacturer’s protocol. We then DNAse treated the RNA (RQ1 RNAse-free DNAse, Promega) according to the manufacturer’s instructions. Finally, we reverse transcribed the RNA using olido dT primers and SuperscriptIII (Thermo Fisher) according to the manufacturer’s instructions.

We performed qPCR using primers 5’-GAC GAC GGC AAC TAC AAG AC-3’ and 5’-TAG TTG TAC TCC AGC TTG TGC-3’ for gfp, primers 5’-GCT TCA AGT AGT CGG GGA TG-3’ and 5’-CCT GTC CCC TCA GTT CAT GT-3’ for mcherry and primers 5’-CGC GAG AAG ATG ACC CAG AT-3’ and 5’-ACA GCC TGG ATA GCA ACG TAC AT-3’ for human β-actin in power SYBRgreen Master mix (Applied Biosystems) according to the manufacturer’s instructions.

We calculated enrichment in qPCR by first calculating the Δct values for IP-Total for a housekeeping gene (β-actin) and the barcode containing transcripts. We then calculated the ΔΔct is by subtracting the actin Δct from the barcode Δct [53].

### Double-IP from neuronal cultures

We applied the cell lysate to 100 μl of anti-HA magnetic beads (Thermo Scientific 88837) that we pre-blocked by overnight incubation with mouse brain lysate. We incubated the lysates on beads at 4^o^C for 8-12 hrs with rotation and washed in PLB3 in ascending salt concentrations as above. We then rinsed the beads in standard PBL3 and eluted the precipitated protein with rotation in 100 μl of 2μg HA peptide (Thermo Fisher 26184) in PBL3 at 37^o^C for 30 min. We collected 10 % of the eluate for Western blot analysis and incubated the rest with pre-blocked anti-FLAG M2 beads (Sigma Aldrich A2220) at 4^o^C for 8-12 hrs with rotation. We washed the beads 6x in PLB3 with ascending salt concentrations as above, rinsed with standard PBL3 and eluted the captured proteins by two sequential incubations with 25 μl of 0.5μg of 3xFLAG peptide (Sigma F4799) in PBL3 at 4^o^C with rotation for 2 hrs. We collected 10 μl of the combined eluate for Western analysis and snapped froze the rest in liquid nitrogen for droplet overlap RT-PCR.

### Western blotting

We used anti-MYC tag antibody, clone A46 (Millipore) and secondary anti-mouse IgG1 HRP-linked antibodies (Cell Signaling 7076s) for the detection of the pre-synaptic proteins and anti-HA tag antibodies, HA.11 (Covance) or anti-HA-HRP 3F10 (Roche 12013819001), for the detection of the post-synaptic proteins. We visualized western blots using either Odyssey fluorescent detection according the manufacturer’s instructions or SuperSignal Western Femto Maximum Sensitivity Substrate (Thermo Scientific 34095).

### HEK cell emulsion RT-PCR

We encapsulated transsynaptic complexes from HEK cells using a custom made setup. We fabricated microfluidic chips as PDMS/glass hybrids using soft-lithography as previously described [54]. We treated the microfluidic channels with a fluorinated tri-chloro silane reagent (heptadecafluoro-1,1,2,2-tetrahydrodecyl) trichlorosilane (Gelest) diluted at 1 % weight in FC3280 oil (3M).

To generate droplets, we actuated the fluids by a set of pressure controllers (MPV1, Proportion Air), which were controlled by a Labview (National Instruments, TX) application via a microprocessor (Arduino). We generated 13 pl (29 μm diameter) droplets at 1 kHz, using a 15 μm deep x 20 μm wide hydrodynamic focusing nozzle. We stabilized the droplets with a PEG-Krytox based surfactant [55] dissolved at 1 % weight in HFE7500 fluorinated oil (3M). Stability through thermocycling was further increased by adding Tetronic 1307 (BASF) at 1 % weight to the RT-PCR mixture [56]. We collected droplets into 0.2 mL PCR tubes and removed the bulk oil phase.

We performed one-step overlap RT-PCR in emulsions as previously described [32]. We made a 100 μL RT-PCR mixture containing: IP products (variable), 25 μl OneStep MasterMix, 5 μl primer 5’-CAG CTC GAC CAG GAT GGG CA-3’ (10 μM), 5 μl Primer 5’-TTC AGC TTG GCG GTC TGG GT-3’ (10 μM), 5 μl primer 5’-TAT TCC CAT GGC GCG CCG CTG GTC GGT ACG GTA ACG GA-3’ (10 μM), 5 μl primer 5’-GGC GCG CCA TGG GAA TAC GGA CGA TGC CGT CCT CGT A-3’ (10 μM), 20 μl 5 % Tetronic, and H_2_O to 100 μl. Briefly, we performed reverse transcription for 30 min at 55^o^C, followed by 2 min at 94^o^C. We preformed PCR amplification with the following thermocycling conditions. 1 cycle of: 94^o^C for 30 sec, 50^o^C for 30 sec, 72^o^C for 2 min; followed by 4 cycles of: 94^o^C for 30 sec, 55^o^C for 30 sec, and 72^o^C for 2 min; followed by 22 cycles of 94^o^C for 30 sec, 60^o^C for 30 sec, 72^o^C for 2 min; followed by a final extension step for 7 min at 72^o^C. After thermal cycling, we visually inspected the emulsion to ensure stability. We broke the emulsions with perfluorooctanol, and collected the aqueous phase for subsequent purification using a spin-column clean-up kit (Promega Wizard SV PCR Cleanup Kit).

We performed a nested PCR amplification (Accuprime Pfx Supermix (Thermo Fisher)) in a total volume of 100 μl using 10 μl of emulsion product as template with 100 nM primers 5’ -AAT GAT ACG GCG ACC ACC GA-3’ and 5’ - CAA GCA GAA GAC GGC ATA CGA-3’ under the following conditions: 2 min 94^o^C; followed by 30 cycles of 94^o^C for 30 sec, 62^o^C for 30 sec, and 72^o^C for 30 sec; followed by a final extension at 72^o^C 7 min.

We purified the final product by gel electrophoresis and sequenced using the PE36 protocol on the MiSeq platform (Illumina, San Diego, CA).

### qPCR to measure crossover rate

We measured the crossover rate in emulsion PCR samples using a quantitative qPCR assay for A-A, B-B, and A-B and B-A barcode pairs. We used primers 5’-CAC ATA AGA CGT GTC CAC CGG TT-3’ and 5’-GGT GCC AGC ATT TTC GGA GGT T-3’ to detect A-A barcode pairs, primers 5’-GCG CAG AGG AAC GCC CAT TTA G-3’ and 5’-GCA GTG GTC GGT GCT CTA AAC T-3’to detect B-B barcode pairs, primers 5’-CAC ATA AGA CGT GTC CAC CGG TT-3’ and 5’-GCA GTG GTC GGT GCT CTA AAC T-3’ to detect A-B barcode pairs and primers 5’-GCG CAG AGG AAC GCC CAT TTA G-3’ and 5’-GGT GCC AGC ATT TTC GGA GGT T-3’ to detect B-A barcode pairs. We performed all qPCR reactions using Power SYBRgreen master mix (Applied Biosystems) and quantified each of the different barcode pairs against a standard.

### Neuron emulsion RT-PCR

To increase throughput of droplet generation, we switched from the custom build setup to a commercially available solution by Dolomite Microfluidics. We produced droplets with 20 μm diameter using a 1R DE Chip with 20 μm etch depth and fluorophilic coating (Dolomite Microfluidics, catalogue number 1864021), pumping fluids using the Mitos P-Pump Remote Basic unit (Dolomite Microfluidics, catalogue number 3200177). We used commercially available oil to encapsulate the droplets (Droplet Generation Oil for probes; Bio-Rad, catalogue number 186-3005) and used the compatible One-Step RT-ddPCR Advanced Kit for Probes (Bio-Rad) with 2 % Tetronic 1307 (BASF) for the emulsion RT-overlap PCR reaction. Per 20 μl RT-PCR rection, we used 4 μl of 10 % Tetronic 1307, 5 μl RT-ddPCR master mix, 2 μl Reverse transcriptase, 1 μl 300mM DTT, 1 μl sample, 1 μl of a 10 μM mixture of 5’-CAG CTC GAC CAG GAT GGG CA-3’ and 5’- TTC AGC TTG GCG GTC TGG GT-3’ and 1 μl of a 1 μM mixture of 5’- TAT TCC CAT GGC GCG CCG CTG GTC GGT ACG GTA ACG GA-3’ and 5’- GGC GCG CCA TGG GAA TAC GGA CGA TGC CGT CCT CGT A-3’.

After droplet generation, we performed reverse transcription at 50^o^C for 60 min, followed by PCR (initial enzyme activation, 95^o^C for 10 min; 40 cycles of 95^o^C for 30 sec, 60^o^C for 1 min; followed by 98^o^C for 10 min). Finally, we broke the emulsions using 3 rounds of phenol-chloroform (Thermo Fisher) cleanup.

We prepared sequencing libraries from the droplet RT-PCR products by performing a nested PCR reaction using Accuprime Pfx Supermix (Thermo Fisher) and primers 5’-AAT GAT ACG GCG ACC ACC GA-3’ and 5’- CAA GCA GAA GAC GGC ATA CGA-3’ at 68^o^C annealing temperature. We gel purified the final product and sequenced it using the PE36 protocol on the MiSeq platform (Illumina).

### Analysis of sequencing data

We error corrected the barcode sequencing reads as previously described [18] and filtered for barcode pairs that contained perfect matches to the zip-codes hardcoded into the barcodes that allowed us to measure crossover rates. We then set an minimum read count for sequences to be analyses to remove template switched and erroneous reads [57]. Finally we calculated the crossover rate of presynaptic barcodes from one experiment joined to postsynaptic barcodes from another, and vice versa, and constructed a connectivity matrix. All sequencing files are accessible on the SRA under accessions SRS1899968 (ZL039: HEK cell emulsions), SRS1899969 (ZL110: neuron emulsions) and SRS1902304 (ZL107: neuron emulsions, high concentration). Code to process these datasets is available in the Supplemental materials.

## Author contributions

I.D.P., J.M.K., D.B., and A.M.Z conceived the study. I.D.P., J.M.K., V.V.V and D.I.R. performed the experiments. I.D.P., E.B. and D.I.R. performed the emulsion experiments. I.R.C. designed and synthesized the bifunctional crosslinkers. J.M.K. and A.M.Z. wrote the paper. A.M.Z. supervised the project.

## Acknowledgements

This work was supported by the following funding sources: National Institutes of Health [5RO1NS073129 to A.M.Z., 5RO1DA036913 to A.M.Z., R01CA181595 to E.B.]; Brain Research Foundation [BRF-SIA-2014-03 to A.M.Z.]; IARPA [MICrONS to A.M.Z.]; Simons Foundation [382793/SIMONS to A.M.Z.]; Paul Allen Distinguished Investigator Award [to A.M.Z.]; PhD fellowship from the Boehringer Ingelheim Fonds to J.M.K.; PhD fellowship from the Genentech Foundation to J.M.K.

## Supplemental Figures

**Supplemental Figure 1.**
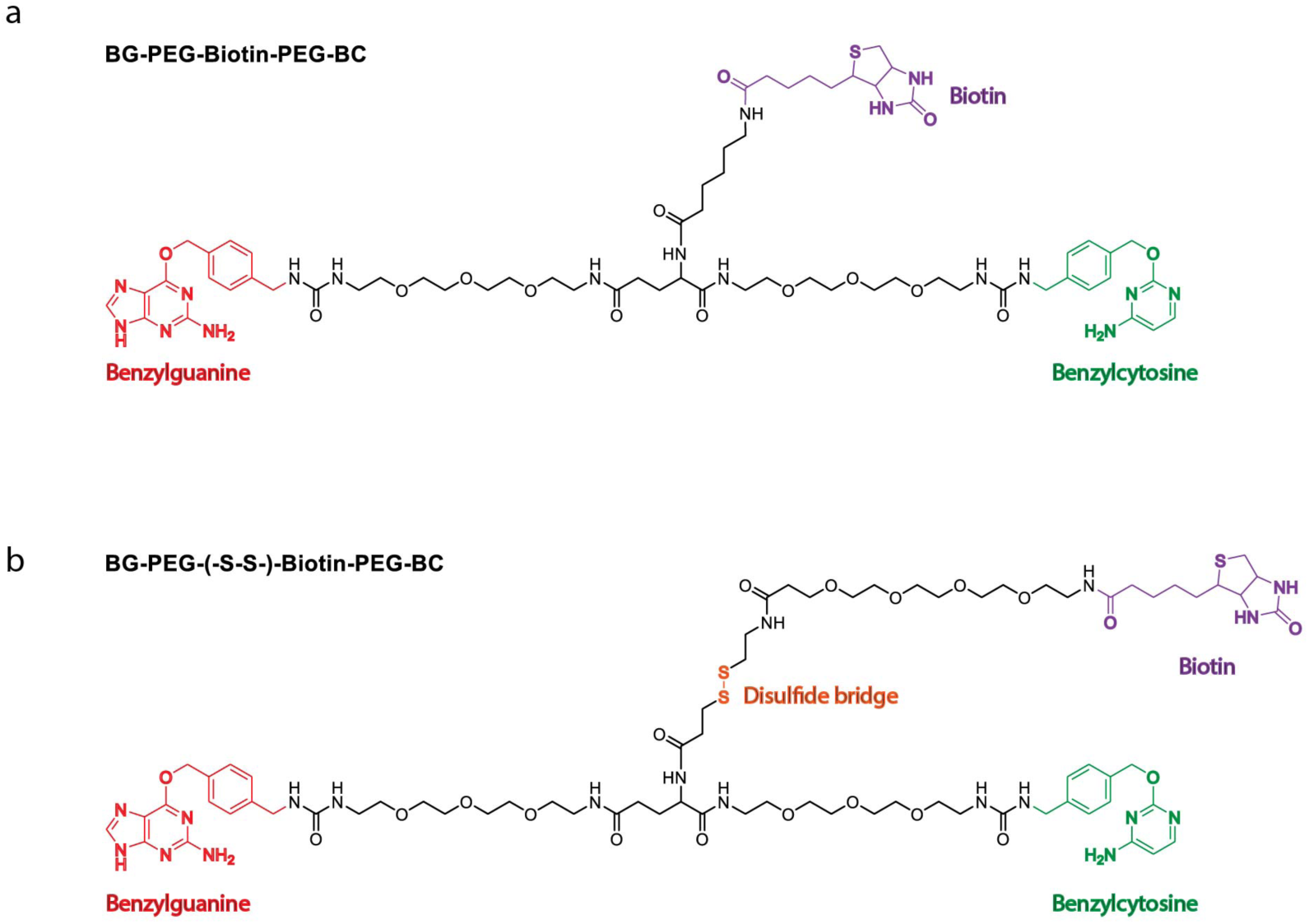
(a) The BG-PEG-Biotin-PEG-BC crosslinker is equipped with functional groups BG and BC, which mediate the covalent tagging of SNAP or CLIP respectively. In addition, the molecule contains a biotin moiety for immunoprecipitation. (b) The BG-PEG-(S-S)-Biotin-BC crosslinker contains a cleavable disulfide bridge for non-denaturing elution in addition to the same functional groups as the BG-PEG-Biotin-BC crosslinker. We use the two crosslinkers interchangeably in this study, unless non-denaturing elution is required.

**Supplemental Figure 2.**
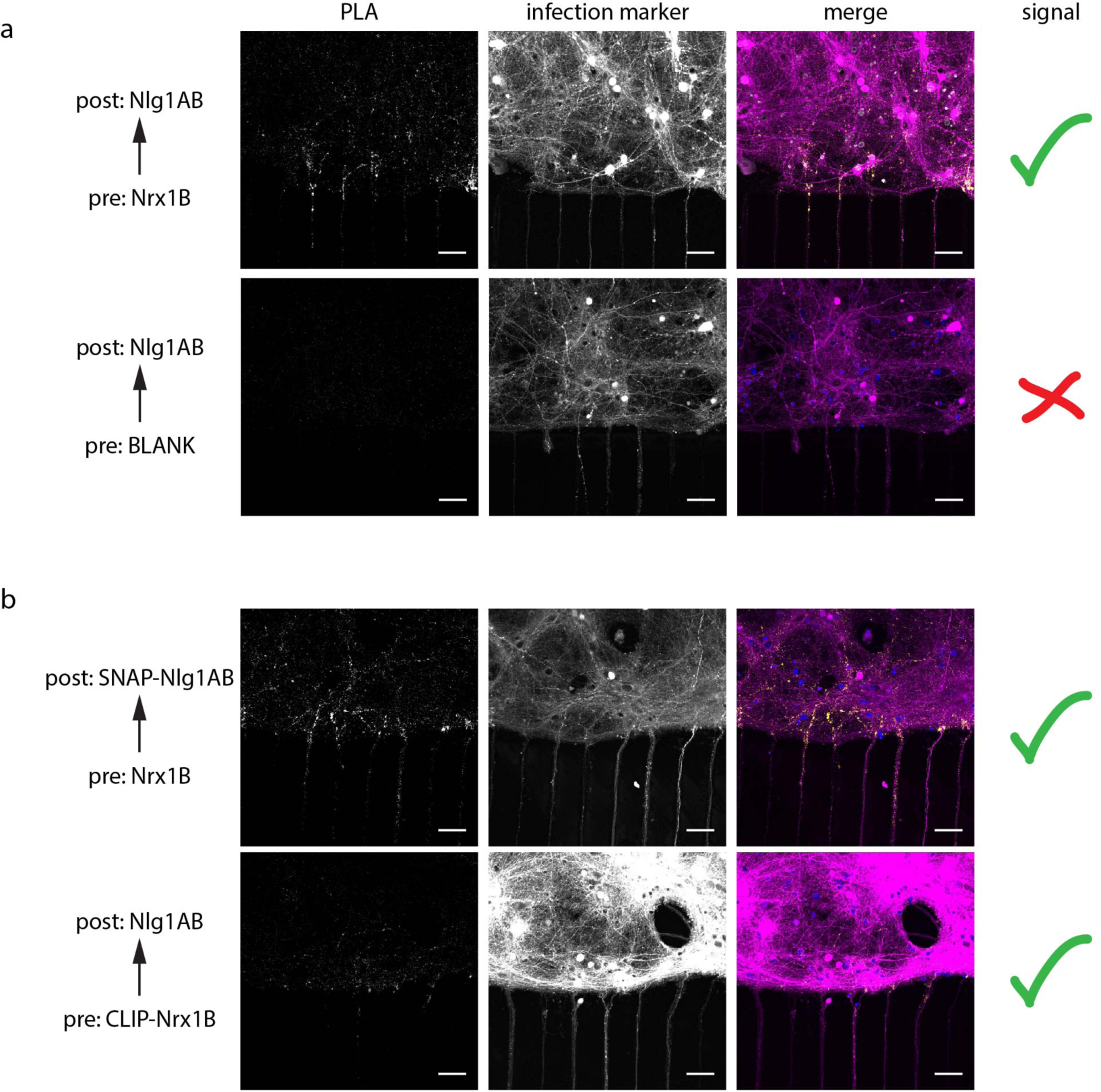
PLA screen for interacting synPRE-P and synPOST-P proteins, part 1. (a) PLA can be used to probe interactions across neurons, for example detecting the interaction of the synaptic proteins MYC-NRXN1B and HA-NLGN1AB. These largely unmodified proteins result in a strong PLA signal across XONAs, acting as a positive control for our screen for synaptically targeted and interacting SYNseq proteins. (b) Addition of CLIP and SNAP domains to the extracellular domains of myc-NRXN1B and HA-NLGN1AB are well tolerated, as indicated by PLA signal across XONAs, when tested together with HA-NLGN1AB or myc-NRXN1B. Scale bar = 50 μm.

**Supplemental Figure 3.**
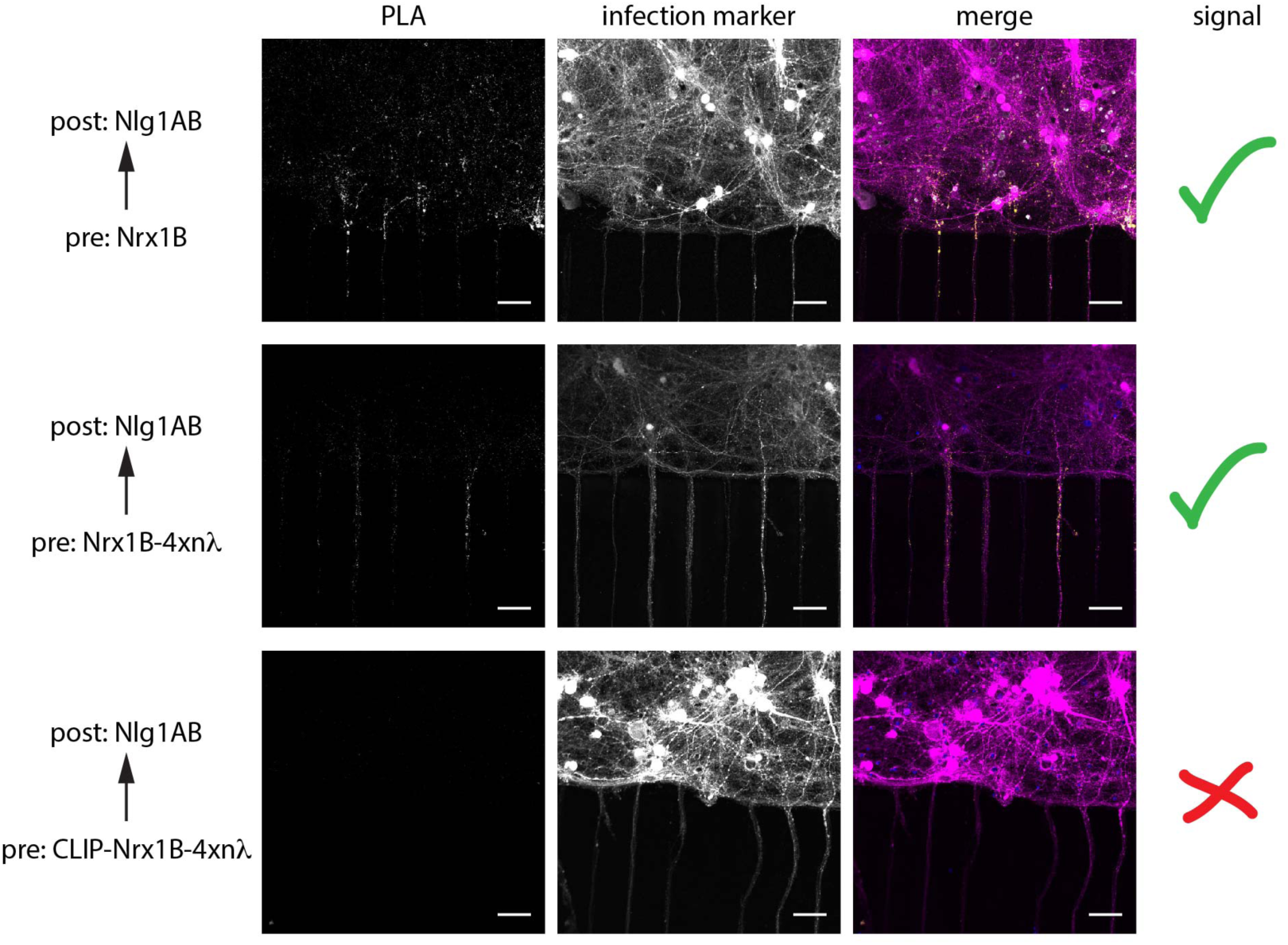
PLA screen for interacting synPRE-P and synPOST-P proteins, part 2. Testing synPRE-P constructs. Addition of four copies of the nλ domain into the c-terminus of myc-NRXN1B is well tolerated (middle) but, surprisingly, PLA signal disappears when we combine CLIP and 4xnλ in a single construct (bottom). Scale bar = 50 μm.

**Supplemental Figure 4.**
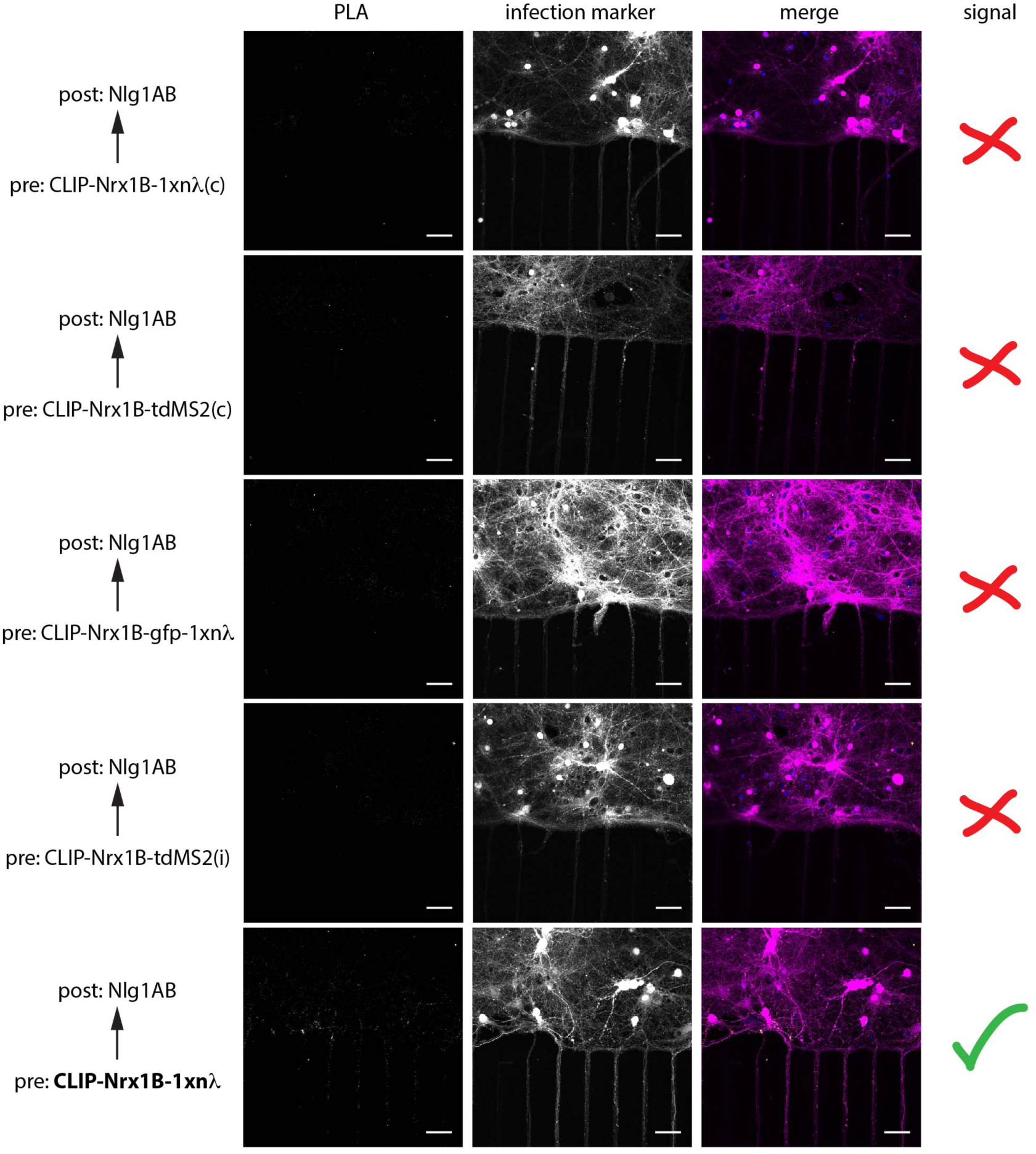
PLA screen for interacting synPRE-P and synPOST-P proteins, part 3. Testing synPRE-P constructs. We tested different positions and spacers for the two RNA binding domains nλ and MS2 in myc-CLIP-NRXN1B. Only a single copy of nλ in the c-terminal tail of myc-CLIP-NRXN1B is tolerated (bottom). Scale bar = 50 μm.

**Supplemental Figure 5.**
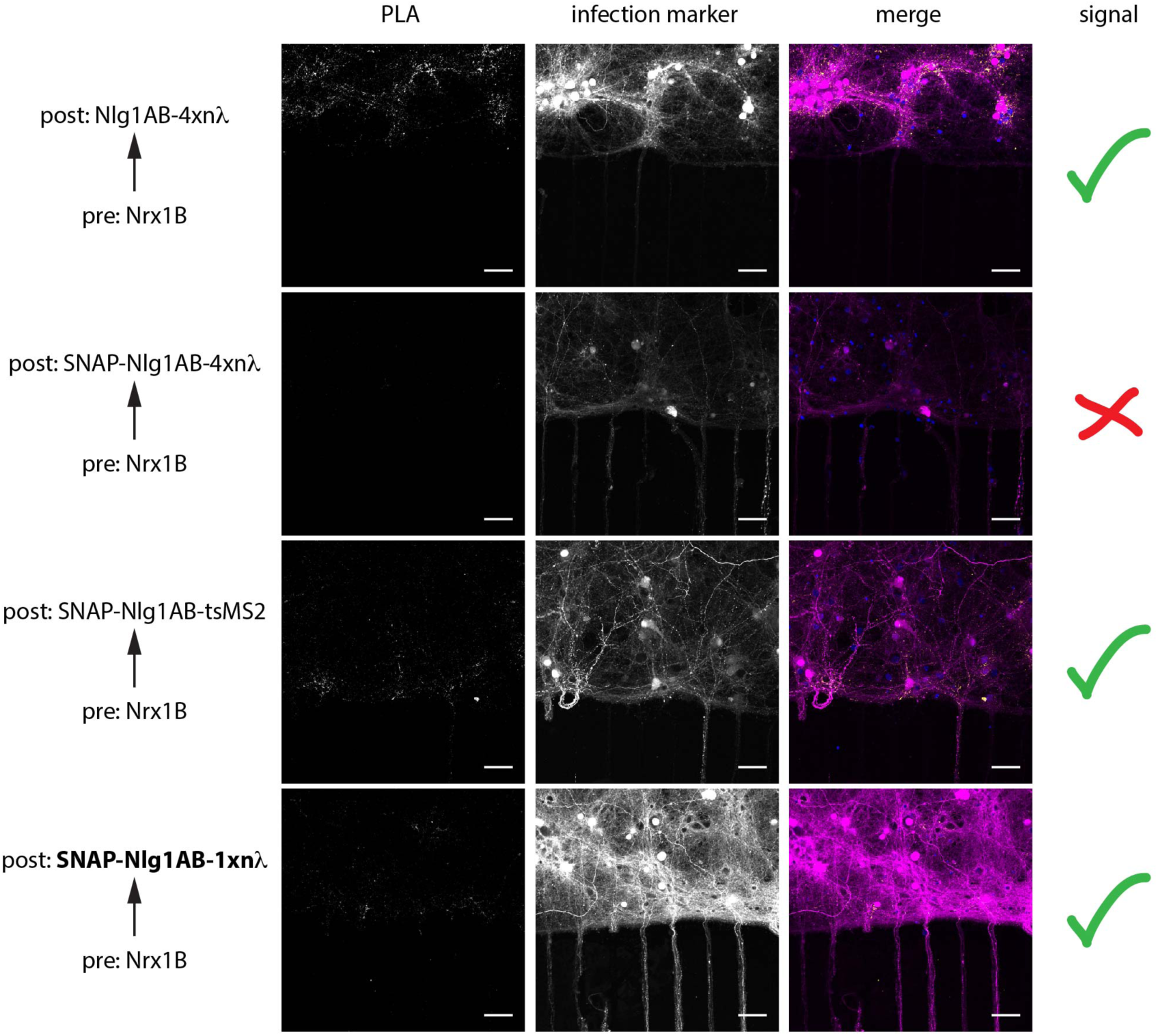
PLA screen for interacting synPRE-P and synPOST-P proteins, part 4. Testing synPOST-P constructs. We tested the RNA binding domains nλ in both four and single copy configurations as well as a tandem dimer of MS2 in HA-SNAP-NLGN1AB. Both insertions of MS2 and of a single copy of nλ in the c-terminal tail are well tolerated. Scale bar = 50 μm.

**Supplemental Figure 6.**
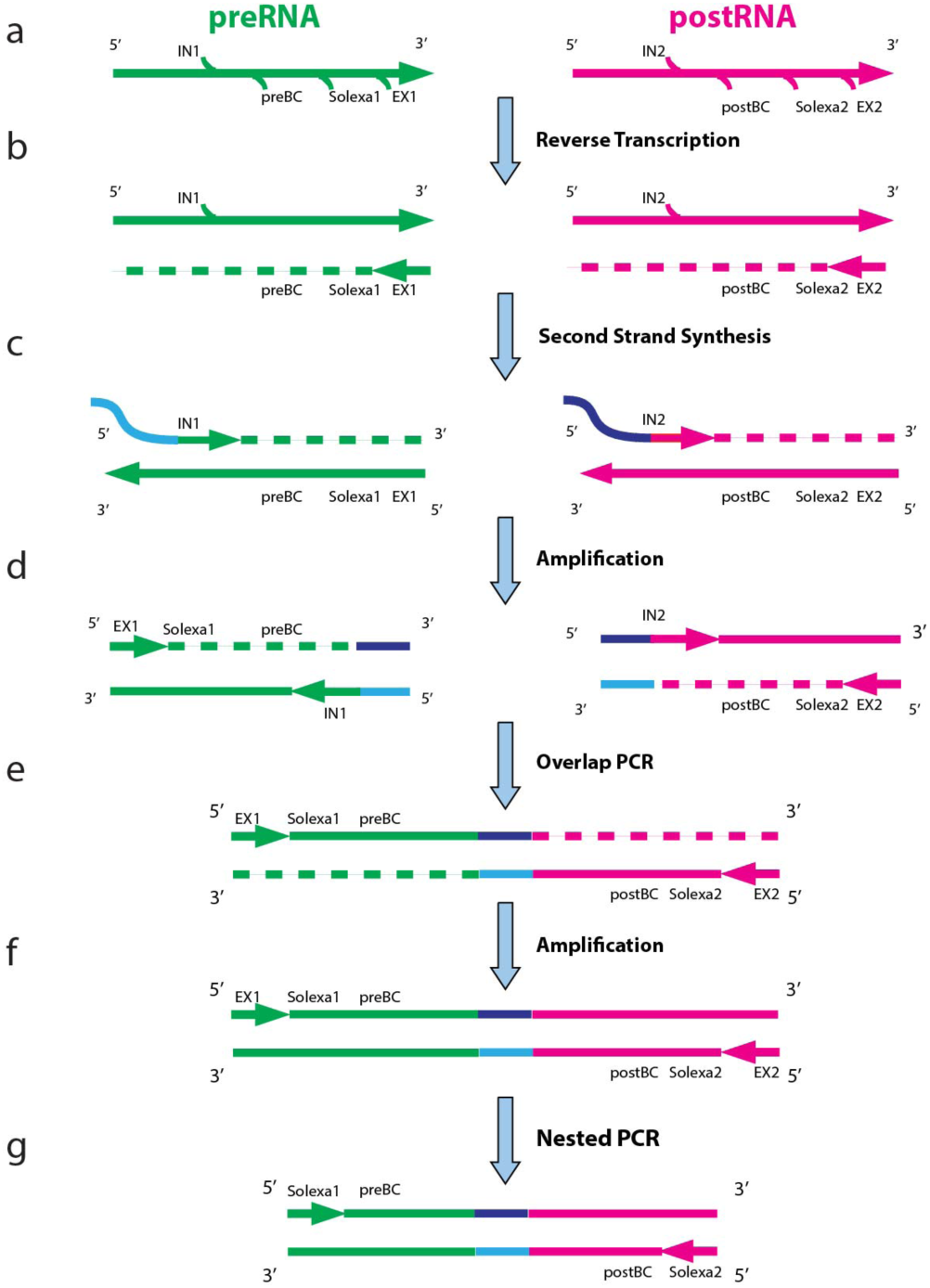
Schematic of overlap RT-PCR. (a) Pre-RNA and Post-RNA molecules containing the barcode sequences, as well as Solexa I and Solexa II sequences for high-throughput sequencing. (b) The RNA molecules are reverse transcribed to cDNA. (c) cDNA is subjected to second strand synthesis using a primer that adds a region of homology between the preRNA and postRNA. (d) Barcodes are amplified individually via an external primer and a limiting concentration of internal primers. (e) Internal primers become limiting and the preRNA and postRNA amplicons perform overlapping PCR. (f) The fused barcode pairs are further amplified using the external primers, before being subjected to (g) nested PCR amplification using the sequencing primers.

**Supplemental Figure 7.**
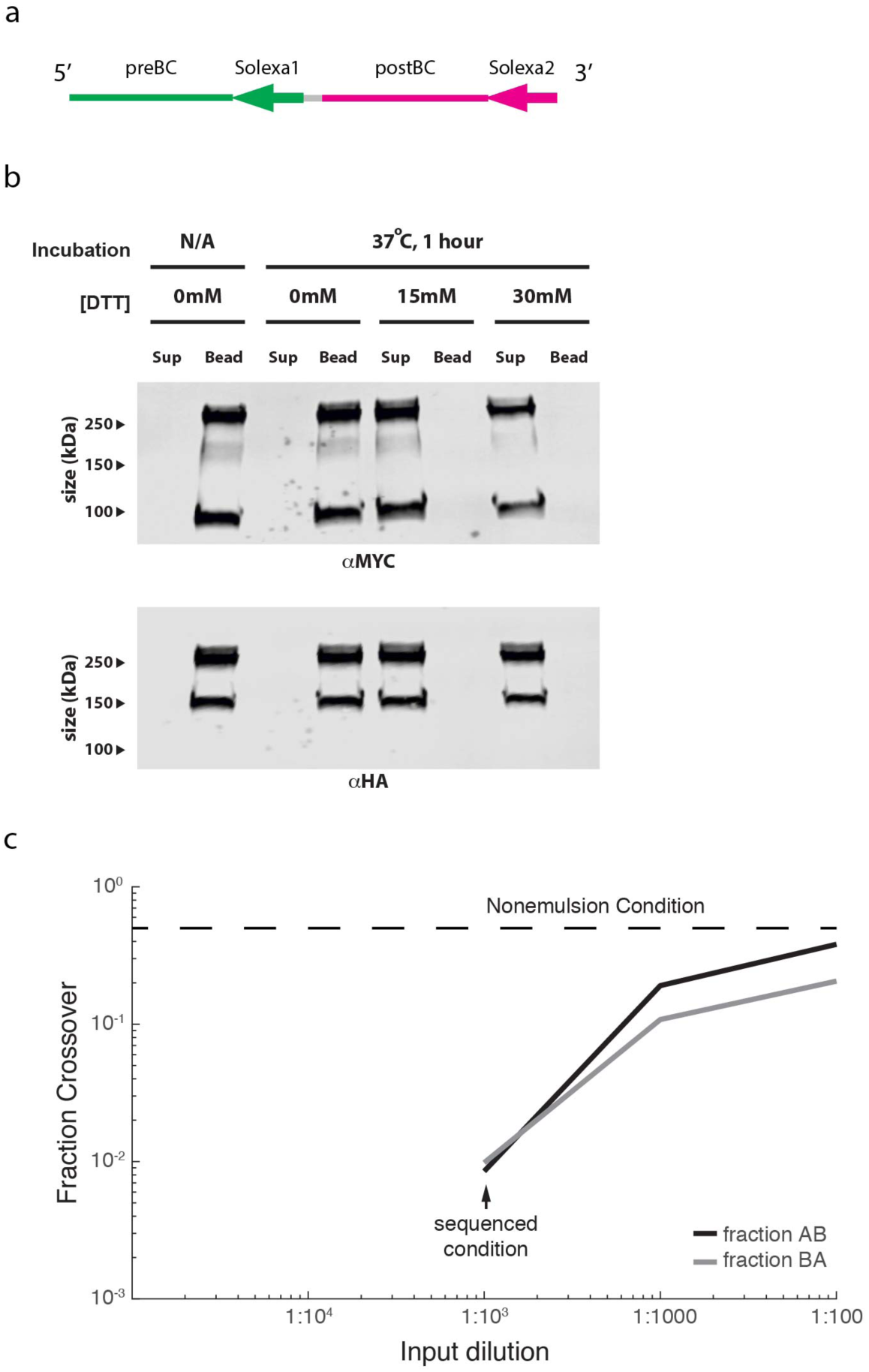
Emulsion RT-PCR from HEK cells. (a) Schematic of the RNA molecules used to benchmark emulsion RT-PCR performance (e.g. in Fig. 6B). Here a single RNA molecule contains both a pre- and post-segment, but in opposite orientations from what is necessary for a sequence-able barcode pair. All the steps described in Supplemental Fig. 6 are therefore necessary to make this RNA into a barcode pair, but no time consuming immunoprecipitations are necessary. (b) Immunoprecipitated complexes from HEK cells co-transfected with synPRE-P and synPOST-P and crosslinked with the BG-PEG-(S-S)-Biotin-PEG-BC crosslinker can be efficiently released from the beads via the application of DTT. (c) qPCR quantification of overlap between the two zip coded experiments in the HEK cell emulsion RT-PCR. We used a dilution of 1:1000 of the IP input for the droplets to create the “connectivity” matrix displayed in Fig. 6C. Further dilution of the IP sample would presumably have resulted in a lower crossover rate, as no asymptotic crossover rate has been reached yet.

**Supplemental Table 1.** Guide to combinations of putatively pre- and postsynaptic proteins tested for interaction across synapses by PLA.

| Presynaptic | Postsynaptic | Result | Figure | Constructs |
| --- | --- | --- | --- | --- |
| Nrx1B | Nlg1AB | Positive | S2 | 56\|57 |
| CLIP-Nrx1B | Nlg1AB | Positive | S2 | 63\|57 |
| Nrx1B-4xλN(i) | Nlg1AB | Positive | S3 | 64\|57 |
| CLIP-Nrx1B-4xλN(i) | Nlg1AB | Negative | S3 | 67\|57 |
| CLIP-Nrx1B-1xλN(c) | Nlg1AB | Weak | S4 | 428\|57 |
| CLIP-Nrx1B-tdMS2(c) | Nlg1AB | Weak | S4 | 430\|57 |
| CLIP-Nrx1B-GFP-1xλN(c) | Nlg1AB | Weak | S4 | 429\|57 |
| CLIP-Nrx1B-tdMS2(i) | Nlg1AB | Weak | S4 | 426\|57 |
| CLIP-Nrx1B-1xλN(i) | Nlg1AB | Positive | S4 | 424\|57 |
| Nrx1B | SNAP-Nlg1AB | Positive | S5 | 56\|65 |
| Nrx1B | SNAP-Nlg1AB-4xλN(i) | Negative | S5 | 56\|68 |
| Nrx1B | SNAP-Nlg1AB-tdMS2(i) | Positive | S5 | 56\|427 |
| Nrx1B | SNAP-Nlg1AB-1xλN(i) | Positive | S5 | 56\|425 |

**Supplemental Table 2.** Constructs used in this study, including Addgene accession numbers.

| Construct name | 1st promoter | 2nd promoter | Express ion | Addgene accession |
| --- | --- | --- | --- | --- |
| **HEK cell characterization** |  |  |  |  |
| IDP431 | CLIP-Nrx1B-4xλN(i) | n/a | HEK | 86999 |
| IDP432 | SNAP-Nlg1AB-1xλN(i) | n/a | HEK | 87000 |
| IDP455 | barcode landing site Gfp-4xboxB | n/a | HEK | 87001 |
| IDP456 | barcode landing site-mCherry-4xboxB | n/a | HEK | 87002 |
| **Neuron characterization** |  |  |  |  |
| jk107 | SNAP-Nlg1AB-1xλN(i) | barcode landing site-mCherry-4xboxB | Sindbis | 87007 |
| IDP466 | myc-3xFlag-CLIP-Nrx1B-1xλN(i) | barcode landing site Gfp-4xboxB | Sindbis | 87008 |
| **HEK cell emulsion experiment** |  |  |  |  |
| IDP253 | MAPP-nλ | n/a | HEK | 87003 |
| IDP254 | postCLAP | n/a | HEK | 87004 |
| IDP378 | preLib-bfp-4xboxB | n/a | HEK | 87005 |
| IDP379 | postLib-mcherry-4xboxB | n/a | HEK | 87006 |
| **PLA screen** |  |  |  |  |
| jk56 | Nrx1B | mCherry | Sindbis | 87009 |
| jk57 | Nlg1AB | mCherry | Sindbis | 87010 |
| jk63 | CLIP-Nrx1B | mCherry | Sindbis | 87011 |
| jk64 | Nrx1B-4xλN(i) | mCherry | Sindbis | 87012 |
| jk67 | CLIP-Nrx1B-4xλN(i) | mCherry | Sindbis | 87013 |
| IDP428 | CLIP-Nrx1B-1xλN(c) | mCherry | Sindbis | 87014 |
| IDP430 | CLIP-Nrx1B-tdMS2(c) | mCherry | Sindbis | 87015 |
| IDP429 | CLIP-Nrx1B-GFP-1xλN(c) | mCherry | Sindbis | 87016 |
| IDP426 | CLIP-Nrx1B-tdMS2(i) | mCherry | Sindbis | 87017 |
| IDP424 | CLIP-Nrx1B-1xλN(i) | mCherry | Sindbis | 87018 |
| jk65 | SNAP-Nlg1AB | mCherry | Sindbis | 87019 |
| jk68 | SNAP-Nlg1AB-4xλN(i) | mCherry | Sindbis | 87020 |
| IDP427 | SNAP-Nlg1AB-tdMS2(i) | mCherry | Sindbis | 87021 |
| IDP425 | SNAP-Nlg1AB-1xλN(i) | mCherry | Sindbis | 87022 |

